# Genomic Insights and Biocontrol Potential of Ten Bacterial Strains from the Tomato Core Microbiome

**DOI:** 10.1101/2024.06.07.597889

**Authors:** Daniele Nicotra, Farideh Ghadamgahi, Samrat Ghosh, Alice Anzalone, Giulio Dimaria, Alexandros Mosca, Maria Elena Massimino, Ramesh Raju Vetukuri, Vittoria Catara

**Affiliations:** Department of Agriculture, Food and Environment, University of Catania, Catania, Italy; Department of Plant Breeding, Swedish University of Agricultural Sciences, Alnarp, Sweden

**Keywords:** Tomato, Microbiome, Endophytes, Rhizosphere, PGPR, BCA, Genomes

## Abstract

Despite their adverse environmental effects, modern agriculture relies heavily on agrochemicals to manage diseases and pests and enhance plant growth and productivity. Some of these functions could instead be fulfilled by endophytes from the plant microbiota, which have diverse activities beneficial for plant growth and health. We therefore used a microbiome-guided top-down approach to select ten bacterial strains from different taxa in the core microbiome of tomato plants in the production chain for evaluation as potential bioinoculants. These taxa included some that are commonly used as biofertilizers and biocontrol agents (*Pseudomonas* and *Bacillus*) as well as the less studied genera *Leclercia, Chryseobacterium, Glutamicibacter,* and *Paenarthorbacter*. When inoculated in the tomato rhizosphere, these strains promoted plant growth and reduced the severity of Fusarium Crown and Root Rot and Bacterial Spot infections. High-quality genomes for each strain were obtained using Oxford Nanopore long-read and Illumina short-read sequencing, enabling the dissection of their genetic makeup to identify phyto-beneficial traits. This yielded a comprehensive inventory of genes from each strain related to processes including colonization, biofertilization, phytohormones, and plant signaling. Traits directly relevant to fertilization including phosphate solubilization and acquisition of nitrogen and iron were also identified. Moreover, the strains carried several functional genes putatively involved in abiotic stress alleviation and biotic stress management, traits that indirectly foster plant health and growth. The gathered genomic information will be instrumental in planning the use of these bacteria individually or in consortia to enhance plant growth by coupling strains with different traits, effects, and mechanisms of action.

## 1 Introduction

Modern farms make extensive use of pesticides to manage diseases and pests as well as chemical fertilizers to enhance productivity (Jacquet et al., 2022). However, the significant negative effects of agrochemicals on both the environment and human health have sparked increasing interest in alternative methods for achieving safe, environmentally sustainable, and eco-friendly crop production (Jacquet et al., 2022). Innovative strategies that minimize reliance on conventional agrochemicals without loss of agricultural productivity and ecological integrity are highly desired.

Current research efforts seeking to reduce agrochemical use are converging towards a prophylactic approach that emphasizes agroecological cropping systems, biodiversity-aware breeding programs, and precision agriculture, with a heavy focus on biological control solutions (Jacquet et al., 2022).

Plant-associated microorganisms play vital roles in protecting plants against abiotic and biotic stress factors (Vessey, 2003; Compant et al., 2005; Singh et al., 2011; Santoyo et al., 2016; Backer et al., 2018; Kelbessa et al., 2023). Some constituents of the plant microbiota actively enhance nutrient uptake, improve nutrient utilization efficiency, and contribute to phytohormone modulation, biocontrol, and the induction of systemic resistance, thereby promoting plant growth and health (Vessey, 2003; Compant et al., 2005; Singh et al., 2011; Pieterse et al., 2014; Santoyo et al., 2016; Backer et al., 2018; Das et al., 2022; Hanifah et al., 2023; Kelbessa et al., 2023). These microorganisms are known as Plant Growth Promoting Microorganisms (PGPM) (Lugtenberg and Kamilova, 2009; Kumar et al., 2022). Endophytes, i.e. microorganisms capable of residing in the internal tissues of host plants, are major constituents of the plant microbiota (Hardoim et al., 2008). Their beneficial effects often exceed those of many rhizosphere-colonizing bacteria and may be especially pronounced when the plant is growing under stress conditions (Hardoim et al., 2008).

Many studies have sought to evaluate the potential of plant-associated microbiomes as PGPM and Biological Control Agents (BCA), but there remains a need to develop diverse biocontrol solutions that can be effectively applied across various environments and management practices (Das et al., 2022; Ayaz et al., 2023). Many Plant Growth-Promoting Rhizobacteria (PGPR) have been identified using bottom-up approaches based on collections of bacteria that display desirable traits in culture-dependent screenings (Compant et al., 2019; Anzalone et al., 2021). However, relying exclusively on culture-dependent selection methods has proven to be a time-intensive strategy that can yield inconsistent results (Berg et al., 2017; Compant et al., 2019).

Research into plant-associated microbial communities has expanded rapidly with the advent of high-throughput sequencing techniques, which have opened up new ways of investigating plant-microbiome and microbe-microbe interactions (Bulgarelli et al., 2012, 2013; Knief, 2014; Du et al., 2020; Kelbessa et al., 2022). For example, top-down approaches were recently used to identify Plant Growth-Promoting (PGP) candidates based on data from microbial community metagenome analyses in an effort to develop biotechnological crop protection strategies (Compant et al., 2019). Microbiome-guided methods for selecting beneficial bacterial strains have mainly focused on the relative abundance and/or enrichment of specific taxa under specific growing conditions (Zhuang et al., 2021), including stress conditions (Kwak et al., 2018; Flemer et al., 2022), or targeting taxa within the ’core’ microbiome (Tian et al., 2017; Bergna et al., 2018; Penyalver et al., 2022).

The core microbiome consists of a set of microbial taxa associated with a specific host or environment along with their genomic and functional characteristics (Lundberg et al., 2012; Neu et al., 2021). It includes microbial taxa that have become vital for plant health as a result of evolutionary processes that have led to the selection and enrichment of taxa that fulfil critical functions for the fitness of the plant holobiont (Lemanceau et al., 2017; Toju et al., 2018; Risely, 2020). Knowledge of core microbiome components, e.g. microbial communities associated with a plant species across various stages of development or under different growing conditions, can thus provide valuable guidance when selecting beneficial microorganisms that could be used to enhance crop resilience and productivity through strategic application of microbial inoculants or other biocontrol agents (Tian et al., 2017; Bergna et al., 2018; Penyalver et al., 2022; Wang et al., 2023).

To plan future tomato microbiome engineering interventions we previously used amplicon-based metagenomics to perform a comprehensive analysis of tomatoes grown under greenhouse conditions spanning the entire plant production chain (Anzalone et al., 2022). The study involved sampling tomato seeds (*Solanum lycopersicum* L. cv. ‘Proxy’) and the rhizosphere of seedlings born from those seeds in a commercial nursery. The development of the seedlings’ root microbiomes was then monitored after transplantation into a greenhouse, and seedling growth in agricultural soil was compared to that on coconut fiber under soilless conditions (Anzalone et al., 2022). The root-associated bacterial communities differed significantly between the nursery and production stages, and also between conventional and soilless conditions in the greenhouse. These findings suggest that it will be essential to account for the variability of the microbiome when seeking to develop biocontrol solutions that will form stable and effective interactions (Anzalone et al., 2022).

The data presented by Anzalone et al. (2022) was used in this work to calculate the core microbiome of the tomato seeds and seedlings at different growth stages under diverse growing conditions to guide the selection of new beneficial bacteria from the same metagenome samples.

Ten bacterial strains from the tomato ‘core microbiome’ collection were characterized and shown to exhibit plant growth promoting and biocontrol properties *in planta*, even though some of them showed no antagonistic activity *in vitro*. Along with strains in the genera *Bacillus* and *Pseudomonas* we shed light onto the PGP phenotypes of strains belonging to the Gram-negative genera *Leclercia* and *Chryseobacterium* and the Gram-positive Micrococcaceae genera *Paenarthrobacter* and *Glutamicibacter*. We show that the construction of high-quality genomes can significantly improve our capacity to investigate and comprehend the intricate mechanisms that make bacterial agents effective in biocontrol.

## 2 Materials and methods

### 2.1 Isolation of bacterial endophytes

The endophytes examined in this study were obtained from seed and root endospheres of tomato (*Solanum lycopersicum* L.) cv. ‘Proxy’ during the course of the metagenomics sample preparation described in Anzalone et al. (2022). More specifically, samples were obtained from the seeds (Seeds_T0) and roots of nursery seedlings before commercialization (Plant_T1_Endo) and from greenhouse tomato plants grown either in agricultural soil (Plant_T2_Soil) or soilless in a coconut fiber substrate (Plant_T2_CF) (Anzalone et al., 2022). Four replicates of 20 seeds and four plant roots bulk samples for each condition were analyzed. Samples were processed according to Anzalone et al. (2021). Cultivable bacterial populations of total, fluorescent, and spore forming bacteria were enumerated in compliance with Anzalone et al. (2021). Bacterial strains were selected using a systematically randomized approach in which solid media plates were divided into six equal parts and colonies from one of the six parts were collected as stated in Bergna et al. (2018). The collected colonies were purified and preserved in 96 microwell cell culture plates (Anzalone et al., 2021).

### 2.2 Molecular and phylogenetic identification of bacteria isolated from the tomato endosphere

The 16S rDNA gene sequence was amplified by PCR and sequenced using the universal 16S rRNA primer pair 27F-1492R (Edwards et al., 1989; Lane, 1991). The master mixtures consisted of 1 x Taq&Go G2 Hot Start colorless PCR Master Mix (Promega), 0.5 μM of each primer, and 1 µL of template in a total volume of 15 μL. Reactions were performed with a GeneAmp® PCR system 9700 thermal cycler using the thermal protocol described by Anzalone et al. (2021). The DNA amplicons were quantified and sequenced by BMR Genomics (Padova, Italy). The nucleotide sequences were searched against the nucleotide collection database of the National Center for Biotechnology Information (NCBI) using the Basic Local Alignment Search Tool BLASTN (http://www.ncbi.nlm.nih.gov). Sequences were aligned using the Clustal-W algorithm as implemented in MEGA XI and deposited in GenBank; the corresponding accession numbers were obtained. A phylogenetic tree was generated based on the alignment profiles using the Neighbor-Joining method (Kumar et al., 2018) with bootstrap-based branch supports in MEGA XI.

### 2.3 Phenotypic characterization of representative bacterial endophytes

*In vitro* tests were conducted as described by Anzalone et al. (2021) to evaluate three plant growth promotion (PGP) traits - siderophore production, phosphate solubilization, and growth on 8% NaCl – in 94 bacterial endophytes from four sample types (Seeds_T0, Plant_T1_Endo, Plant_T2_Soil, Plant_T2_CF). The production of hydrogen cyanide (HCN) and 1-aminocyclopropane-1-carboxylic acid (ACC) deaminase was assessed using the methods of Strano et al. (2017) and Penrose and Glick, (2003), respectively. All experiments were performed in three independent replicates.

### 2.4 Antimicrobial activity of representative bacterial endophytes

Bacterial endophytes were tested on PDA plates for *in vitro* antagonistic activity according to Anzalone et al. (2021), against the following pathogens: the bacteria *Clavibacter michiganensis* subsp*. michiganensis* strain PVCT 156.1.1 (Cmm), *Pseudomonas syringae* pv. *tomato* strain PVCT 28.3.1 (Psto), *Xanthomonas euvesicatoria* pv. *perforans* strain NCPPB4321 (Xep) and the fungi *Fusarium oxysporum* f. sp*. radicis-lycopersici* strain PVCT127 (Forl), and *Botrytis cinerea* strain Bc5 (Bot). Briefly, bacterial pathogen suspensions were normalized to an OD_600_ of 0.1 (Anzalone et al., 2021) and inhibition halo radii (in mm) were measured after 48h of incubation. For fungal pathogens a mycelial plug was placed in the center of the plate (Anzalone et al., 2021) and the antifungal activity was expressed as a Percentage of Growth Inhibition (PGI) according to Vincent (1947). All strains were tested in three independent replicates.

### 2.5 Selection of strains for further trials

To identify key bacterial components of the tomato plant-associated samples, the core microbiome of the samples reported by Anzalone et al. (2022) was calculated. The samples used in the analysis represented tomato seeds (Seed_T0), the root rhizosphere (Plant_T1_Rhizo) and endorhizosphere (Plant_T1_Endo) of tomato plants ready for sale; and the rhizosphere and endorhizosphere of tomato plants at flowering and fruit set after transplantation into agricultural soil (Plant_T2_Soil_Rhizo; Plant_T2_Soil_Endo) and coconut fiber bags (Plant_T2_CF_Rhizo; Plant_T2_CF _Endo). Following the selection criteria of Hamonts et al. (2018), bacterial core taxa up to the genus level with prevalences ≥ 75% in the aforementioned samples were investigated using the microbiome package in R (Shetty and Lahti, 2019). Strains from the established collection were chosen for further trials based on core microbiome analysis and growth stability *in vitro*.

### 2.6 *In planta* bioassays

#### 2.6.1 Microorganisms’ growing conditions and inoculum preparation

Bacterial strains were grown on NDA plates for 24 h at 27 ± 1 °C. Single colonies were inoculated in 25 mL of LB broth and incubated for 24 h at 27 ± 1 °C in a rotary shaker (180 rpm). The bacterial cultures were centrifuged at 5,000 rpm for 15 min, and after discarding the supernatant, the pellets containing the bacterial cells were resuspended in sterile water and the density was adjusted to 1 · 10^8^ colony forming units (cfu) · mL^-1^.

Xep suspensions were prepared as above, with a final concentration of 1 · 10^8^ cfu · mL^-1^ (Anzalone et al., 2021). To produce Forl inoculum, fresh conidia were collected from sporulating colonies grown for 14 days on PDA at 23 °C. Petri dishes were flooded with 10 ml of sterile distilled water, then conidia were scraped using sterile spatulas and transferred to sterile 50 ml tubes. After filtration through four layers of cheesecloth, the concentration of the resulting spore suspension was estimated using a hemocytometer under light microscopy and adjusted to 4 · 10^6^ conidia · mL^-1^ (Manzo et al., 2016).

#### 2.6.2 Plant material and growing conditions

Tomato plantlets of the variety Moneymaker were produced from seeds in growth chamber. Briefly, seeds were surface-sterilized by immersion in 3% sodium hypoclorite for 5 min followed by three washing steps in sterile water (Ghadamgahi et al., 2022) and dried on sterile filter paper in a laminar flow cabinet. Seed were sown in trays filled with a commercial potting substrate (Krukväxtjord Lera/Kisel, SW Horto). Trays were covered with plastic bags and kept in a growth chamber under controlled conditions (22 °C/16 h light, 18 °C/8 h dark, 60% relative humidity). After germination, the chamber conditions were changed to 25°C:22°C day:night. The light intensity was set at 300 µmol · m^-2^s^-1^ (Fan et al., 2013) and then changed to 225 µmol m^-2^s^-1^ when the plants were three weeks old. Plants were transplanted into 2 L volume pots for subsequent experiments.

#### 2.6.3 *In vivo* biocontrol activity

The biocontrol activity of the selected bacterial endophytes was assessed against the causal agent of Tomato Crown and Root Rot, *Fusarium oxysporum* f. sp. *radicis-lycopersici* (strain PVCT127), and *Xanthomonas euvesicatoria* pv. *perforans* (strain NCPPB4321) one of the causal agents of Tomato Bacterial Spot (Osdaghi et al., 2021). Six plants were used as replicates in each pathogen/endophyte combination. Endophytes were applied by seed soaking (30 minutes) and soil drenching with 50 mL of the bacterial endophyte suspension, three weeks after plant emergence and after approximately three further weeks but exactly 72 or 24 h before *Fusarium* and *Xanthomonas* inoculation, respectively.

For the artificial inoculations with Forl, 30 ml of conidial suspension was poured into the soil near each tomato plant and a wound in the crown was made by a razor blade to assist pathogen penetration. Control plants were wounded in the same way but inoculated with sterile water. The growth chamber was set at 22 °C/16 h light and 20 °C/18 h dark, with 80% relative humidity. Disease evaluation was performed 45 days after Forl inoculation (Vitale et al., 2014). All seedlings were gently uprooted and their crowns and stems were examined. To determine disease incidence, all plants were sectioned to ascertain the presence of disease symptoms and the percentage of infected tomato plants was determined. Disease severity was assessed by measuring the length (cm) of vascular discoloration in each tomato stem.

For the bacterial spot biocontrol assay, Xep cell suspensions (1 · 10^8^ cfu · mL^-1^) or water as a negative control, supplemented with 0.01% tween 20, were spray-inoculated on tomato plants. Plants were covered with plastic bags 24 h before pathogen inoculation and remained covered for the following 72 h to maintain a relative humidity of around 100%. The growth chamber was set at 26 °C/16 h light and 24 °C/18 h dark, with 80% relative humidity. Six days after inoculation, the disease incidence was recorded and the disease severity was calculated by estimating the percentage of the leaf area affected (necrotic tissue) by bacterial spot in approximately 10 leaflets using the ImageJ software (https://imagej.nih.gov/ij/).

#### 2.6.4 Tomato growth promotion assay

Growth promotion activity was evaluated using a completely randomized block experimental design. After transplanting, 20 mL of the appropriate bacterial suspension (or water as a negative control) was added to each pot by soil drenching (Anzalone et al., 2021). Pots were observed regularly and watered daily as needed. Shoot height was recorded at five different time points: T0 (treatment) and T1-4 (from 1 to 4 weeks after treatment). After one month, the seedlings were uprooted and the fresh and dry shoot and root weights were determined. For dry weight measurements, plant shoots and roots were oven-dried at 70 °C for three days before weighing. Seven replicates were used for each treatment.

### 2.7 DNA extraction and whole genome sequencing

Bacterial strains were grown in LB broth inoculated with a single bacterial colony from a 24-h-old culture on NDA and incubated overnight at 27±1°C under continuous shaking (180 rpm). Total genomic DNA was extracted from bacterial cultures using the Wizard® HMW DNA Extraction Kit (Promega) according to the manufacturer’s instructions. Complete bacterial genome sequences were determined by a combination of long and short reads. Long and short read sequencing were performed with an Oxford Nanopore GridION X5 platform and an Illumina NovaSeq 6000 platform (paired-end read length, 150 bp), respectively.

### 2.8 Pre-processing of reads, genome assembly and annotation

The quality of raw Nanopore reads was checked with NanoPlot v1.42.0 (De Coster and Rademakers, 2023). Adapters were trimmed with Porechop_ABIv0.5.0 (Bonenfant et al., 2022). Seqkit v2.8.1(Shen et al., 2016) was used for quality filtering with 1,000 bp read length and Q10 quality cutoffs. Filtered nanopore reads were assembled using Flye v.2.9.4 (Kolmogorov et al., 2019). PILON v1.24 (Walker et al., 2014) was used for polishing with Illumina reads.

Illumina raw reads were pre-processed (adapter trimming, quality filtering [>Q30] and quality checking) with fastp v 0.23.4 (Chen et al., 2018). Filtered reads were fed into SPAdes v3.15.5 (Bankevich et al., 2012) for assembly.

CheckM v1.1.6 (Parks et al., 2015) was used to determine the completeness and contamination of the assemblies. Assembly statistics were computed with QUAST v5.2.0 (Gurevich et al., 2013). Plasmer (Zhu et al., 2023) was used to identify plasmid sequences. General annotation of genomes was performed using Prokka v1.14.5 (Seemann, 2014).

GTDB-Tk v2.3.0 (Chaumeil et al., 2022) was used for taxonomic annotation of each genome. Genome sequence data were uploaded to the Type (Strain) Genome Server (TYGS), a free bioinformatics platform available at https://tygs.dsmz.de, to perform whole genome-based taxonomic analyses (Meier-Kolthoff and Göker, 2019). Genomic relatedness was determined using average nucleotide identity (ANI) values computed with EzBioCloud (Yoon et al., 2017). Two genomes belonging to the same species should have a dDDH of at least 70%, corresponding to an ANI of at least 95% (Goris et al., 2007; Auch et al., 2010; Meier-Kolthoff et al., 2013). Plant growth promoting traits (PGPT) were predicted using the PGPT-Pred module of PLaBAse v1.01 (Patz et al., 2021). PIFAR-BASE was used to identify ‘plant bacterial only interaction factors’ from the annotated protein files for each strain using the BlastP+HMMER Aligner/Mapper (Patz et al., 2021). The bacterial version of antiSMASH 7.0 (Blin et al., 2023) was used to screen for secondary metabolites.

### 2.9 Statistical analysis

Data from the PGP and biocontrol experiments were analyzed by analysis of variance (ANOVA) using Minitab 20 statistical software (Minitab, Inc., State College, PA). Means were separated using Tukey’s post-hoc HSD test.

### 2.10 Data availability

16S rRNA gene sequences of the strains used in this work were submitted to the GenBank database under accession numbers from MZ066824 to MZ066917.

All of the assembled genomes and respective raw reads are available under BioProject ID: PRJNA1096641.

## 3 Results

### 3.1 Isolation and identification of bacterial endophytes

The bacterial endophytes examined in this study were obtained from samples that were prepared for metagenomic analysis of microbial communities in tomato plants at multiple stages in the cultivation chain from nursery to greenhouse (Anzalone et al., 2022). Bacteria (total, fluorescent, and spore-forming) were enumerated on different media from samples obtained from the endospheric compartments of tomato cv. ‘Proxy’ seeds (Seeds_T0), from the roots of nursery-grown seedlings at the commercialization stage (Plant_T1_Endo), and from the roots of two-month-old plants that had been transplanted into agricultural soil (Plant_T2_Soil_Endo) or coconut fiber bags (Plant_T2_CF_Endo). The total, spore forming, and fluorescent bacterial population sizes in the seeds were 1.46, 0.4, and 0.8 log CFU per gram of seed, respectively. The root endosphere bacterial concentrations of adult plants grown in agricultural soil and coconut fiber substrate were similar, and both were higher than the bacterial titers of plantlets in the nursery (Supplementary Figure S1).

Ninety-four representative bacteria from the endospheric (root or seed) compartments were selected for further investigation from a collection of approximately 2000 bacterial colonies identified across the entire experiment.

BLASTN similarity matches of the 16S rRNA genes sequence indicated that the selected strains belonged to genera from seven orders (Supplementary Table S1): the Gram-positive Bacillales and Micrococcales, and the Gram-negative Pseudomonadales, Enterobacteriales, Flavobacteriales, Burkholderiales, and Xanthomonadales. More specifically, the Bacillales strains belonged to the genera *Bacillus*, *Paenibacillus*, *Staphylococcus*, and *Priestia*; the Micrococcales strains belonged to the genera *Glutamicibacter*, *Microbacterium*, *Curtobacterium*, *Paenarthrobacter*, and *Arthrobacter*; the Pseudomonadales strains belonged to the genus *Pseudomonas*; the Enterobacteriales strains belonged to *Enterobacter*, *Ewingella*, and *Serratia*; Flavobacteriales was represented by the genera *Flavobacterium* and *Chryseobacterium*; Burkholderiales was represented by a single strain of the genus *Delftia*; and Xanthomonadales was represented by multiple strains in the genus *Stenotrophomonas*. Sequences were deposited at GenBank under accession numbers from MZ066824 to MZ066917 (Supplementary Table S1). Measurements of the relative abundance of the cultivable bacteria at the order level in the different endosphere samples showed that the majority of the strains belonged to the order Bacillales and Pseudomonadales (Figure 1A). A dendrogram showing the phylogenetic relationships of the selected bacterial strains is shown in Figure 1B.

**Figure 1.**
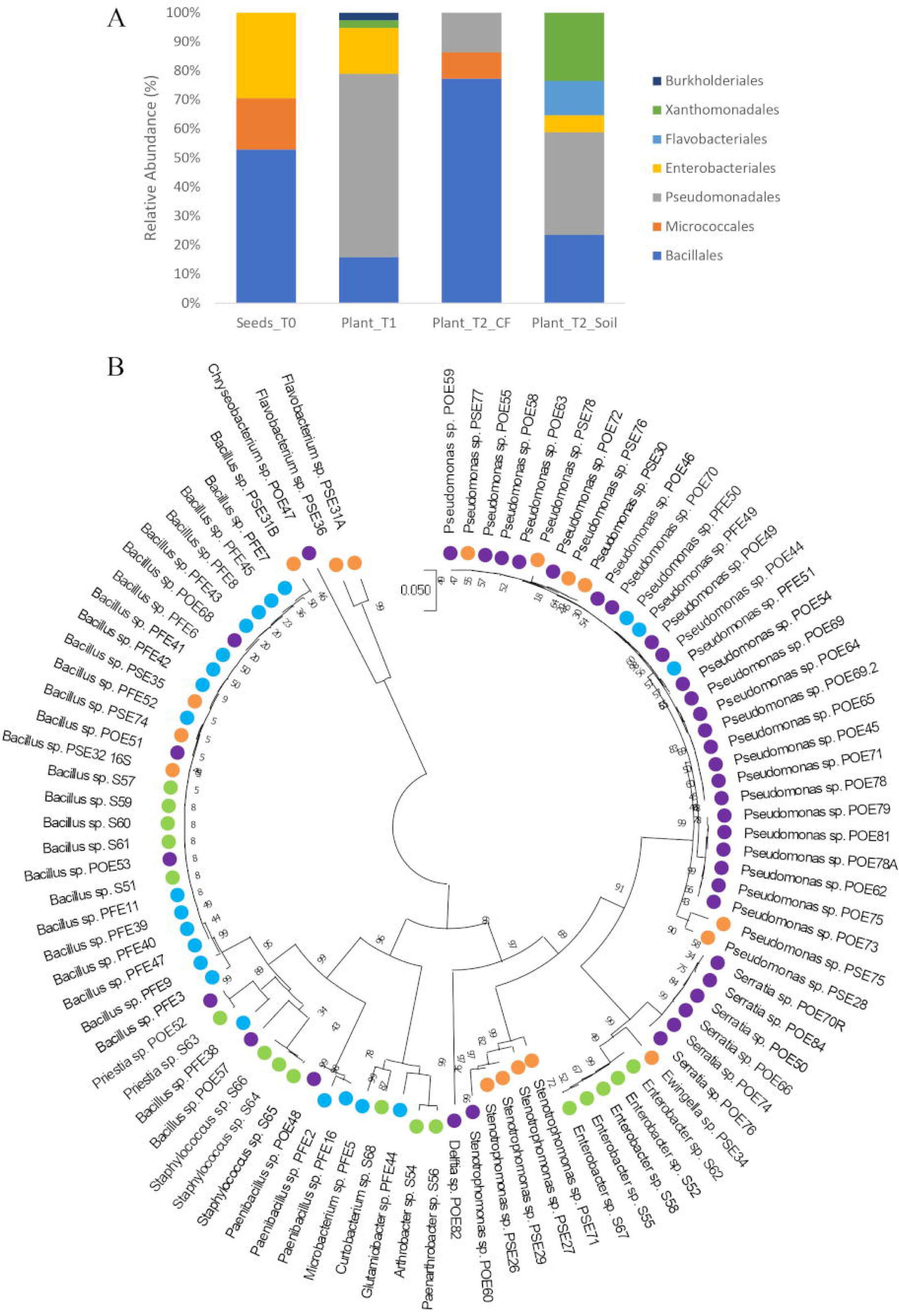
(A) Distribution of cultivable bacterial communities in the endosphere samples of tomato seeds (Seeds_T0) and roots (Plant_T1, Plant_T2_CF, and Plant_T2_Soil) at the taxonomic order level; (B) Phylogenetic tree based on 16S rRNA gene sequences of the 94 endophytic strains isolated in this study. The evolutionary history was inferred using the Neighbor-Joining method. The evolutionary distances were computed using the Tamura 3-parameter method. There was a total of 731 positions in the final dataset. Colors highlight the isolation source of each strain: green, Seeds_T0; violet, Plant_T1; blu, Plant_T2_CF; orange, Plant_T2_Soil.

### 3.2 Phenotyping of beneficial bacterial traits

The bacterial strains were characterized for different beneficial properties, revealing that a high percentage of strains showed PGP traits. Approximately 87% of the bacterial strains from the tomato endospheric compartments could grow in 8% NaCl, while 51% were able to produce siderophores and solubilize insoluble organic phosphate. However, only 2% and 21% of the strains were positive for HCN and ACC deaminase production, respectively (Supplementary Figure S2A). Approximately 30% of the bacteria (28 out of 94 strains) showed antagonistic activity towards all of the tested phytopathogenic bacteria and fungi (Supplementary Figure S2B), while around 45% of the strains (41 out of 94) were antagonistic to all the bacterial pathogens. The highest antimicrobial activity (based on the number of antagonistic strains and inhibition zone radius) was observed against *C. michiganensis* subsp. *michiganensis* PVCT 156.1.1, followed by *P. syringae* pv. *tomato* PVCT 28.3.1. Roughly 54% of the endophytic strains (51 out of 94) inhibited the mycelial growth of the fungal targets *Fusarium oxysporum* f. sp. *radicis-lycopersici* PVCT127 and *Botritys cinerea* Bc5 to at least some degree when compared to a non-challenged colony. Moreover, 26% of the strains achieved at least 60% growth inhibition against the former fungal pathogen, while 11% of the strains achieved the same level of inhibition against the latter (Supplementary Figure S2B).

### 3.3 Selection of strains from the core microbiome for further trials

The analysis of the bacterial communities in the tomato samples made it possible to identify twenty-seven core microbiome genera (i.e., genera with a prevalence of at least 75%) in the tomato seed, rhizosphere, and endorhizosphere samples from the experimental trials of Anzalone et al. (2022) (Figure 2). *Flavobacterium*, *Pseudomonas* and *Bacillus* were the most abundant core microbiome genera. Seven of the core genera were also represented in bacterial strains obtained from the endosphere samples by *in vitro* culturing, namely *Flavobacterium*, *Pseudomonas*, *Bacillus*, *Enterobacter*, *Chryseobacterium*, *Arthrobacter* and *Stenotrophomonas*. At least one strain from each of these genera was selected for further investigation (Table 1); in the cases of *Pseudomonas* and *Bacillus*, two and three representatives were chosen, respectively (Table 1). *Flavobacterium* and *Stenotrophomonas* strains were not selected due to their low growth stability *in vitro*. BLASTN comparisons of the 16S rRNA gene sequences of these bacterial strains in axenic culture against the sequences of the core OTUs revealed putative matches for the strains *Arthrobacter* S54, *Paenarthrobacter* S56, *Glutamicibacter* PFE44 and *Pseudomonas* POE54 and POE78A (data not shown).

**Figure 2.**
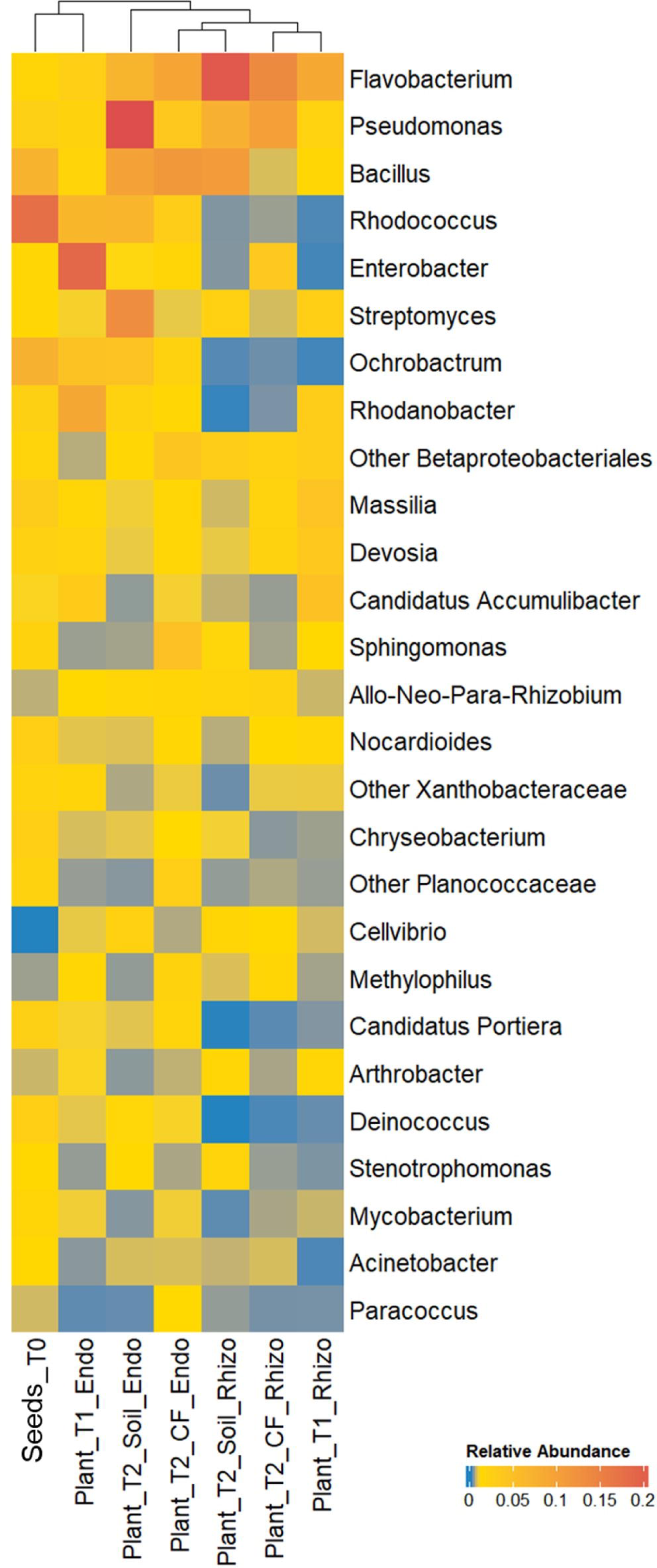
Bacterial genera of the tomato core microbiome showing a prevalence ≥ 75% in the samples of the bacterial communities in the tomato growth chain according to Anzalone et al. (2022). Bacterial genera are indicated in the right; samples are indicated on the bottom (see Material and Methods section for sample details).

**Table 1.** Isolation source and identification by partial 16S rRNA gene and whole genome sequencing of the ten bacterial endophytes selected based on the core microbiome analysis.

| Strain ID | Isolation source<br>(substrate) | Class | Order | Family | Identification |  |
| --- | --- | --- | --- | --- | --- | --- |
|  |  |  |  |  | 16S rRNA gene <sup>a</sup> | Whole genome <sup>b</sup> |
| POE54 | Roots (peat) | Gammaproteobacteria | Pseudomonadales | Pseudomonadaceae | *Pseudomonas fluorescens* | *Pseudomonas salmasensis* |
| POE78A | Roots (peat) | Gammaproteobacteria | Pseudomonadales | Pseudomonadaceae | *Pseudomonas extremorientalis* | *Pseudomonas simiae* |
| POE47 | Roots (peat) | Flavobacteria | Flavobacteriales | Weksellaceae | *Chryseobacterium* sp. | *Chryseobacterium* sp. |
| S52 | Seeds | Gammaproteobacteria | Enterobacteriales | Enterobacteriaceae | *Enterobacter* sp. | *Leclercia* sp. |
| PSE31B | Roots (soil) | Bacilli | Bacillales | Bacillaceae | *Bacillus amyloliquefaciens* | *Bacillus velezensis* |
| PFE42 | Roots (coconut fiber) | Bacilli | Bacillales | Bacillaceae | *Bacillus velezensis* | *Bacillus velezensis* |
| PFE11 | Roots (coconut fiber) | Bacilli | Bacillales | Bacillaceae | *Bacillus subtilis* | *Bacillus velezensis* |
| PFE44 | Roots (coconut fiber) | Actinomycetes | Micrococcales | Micrococcaceae | *Glutamicibacter halophytocola* | *Glutamicibacter halophytocola* |
| S54 | Seeds | Actinomycetes | Micrococcales | Micrococcaceae | *Arthrobacter* sp. | *Paenarthrobacter ureafaciens* |
| S56 | Seeds | Actinomycetes | Micrococcales | Micrococcaceae | *Paenarthrobacter* sp. | *Paenarthrobacter* sp. |

<sup>a</sup> The nucleotide sequences were searched against the nucleotide collection database at the National Center for Biotechnology Information (NCB nucleotide database using Basic Local Alignment Search Tool BLASTN (http://www.ncbi.nlm.nih.gov). <sup>b</sup> Complete bacterial genome sequences were determined by a combination of long and short reads. For long and short reads sequencing, an Oxford Nanopore GridION X5 platform and Illumina NovaSeq 6000 platform were used, respectively.

### 3.4 Genome sequencing of beneficial endophytes

The genetic potential of the selected beneficial bacteria was explored by using a combination of long-read Oxford Nanopore and short-read Illumina sequencing to obtain their genomes in order to identify traits associated with plant growth promotion and biocontrol. The genomes were *de novo* assembled to create high quality, complete reference genomes, yielding individual genomes whose characteristics are detailed in Table 2. Final assembly quality was validated by estimating contamination and completeness with CheckM (Parks et al., 2015; Table 2). Overall, the assembled genomes showed over 98% completeness and <1% contamination (Table 2).

**Table 2.** General features of the genomes of the 10 strains selected.

|  | Phylogenetic<br>affiliation <sup>a</sup> | Completeness<br>(%) | Contamination | Size<br>(Mbp) | N<sub>50</sub> | L<sub>50</sub> | GC<br>(%) | Genes | CDS | rRNA | tRNA | mis<br>RN |
| --- | --- | --- | --- | --- | --- | --- | --- | --- | --- | --- | --- | --- |
| POE54 | *Pseudomonas*<br>*salmasensis* | 98.99 | 0.40 | 6.08 | 6.09 | 1 | 60.31 | 5678 | 5502 | 19 | 73 | 85 |
| POE78A | *Pseudomonas simiae* | 99.51 | 0.32 | 6.22 | 6.22 | 1 | 60.33 | 5995 | 5838 | 19 | 75 | 62 |
| POE47 | *Chryseobacterium* sp. | 98.53 | 0.25 | 4.39 | 4.40 | 1 | 36.31 | 4481 | 4388 | 15 | 63 | 14 |
| S52 | *Leclercia* sp. | 99.28 | 0.26 | 4.93 | 4.93 | 1 | 56.78 | 5052 | 4820 | 25 | 86 | 12 |
| PSE31B | *Bacillus velezensis* | 98.79 | 0.00 | 4.02 | 4.02 | 1 | 46.41 | 4374 | 4178 | 27 | 87 | 81 |
| PFE42 | *Bacillus velezensis* | 99.42 | 0.25 | 4.21 | 2.49 | 1 | 45.95 | 4534 | 4333 | 28 | 90 | 82 |
| PFE11 | *Bacillus velezensis* | 96.91 | 0.54 | 3.99 | 3.99 | 1 | 46.41 | 5210 | 5017 | 27 | 86 | 80 |
| PFE44 | *Glutamicibacter*<br>*halophytocola* | 93.73 | 0.92 | 3.72 | 3.72 | 1 | 60.40 | 4194 | 4086 | 19 | 65 | 25 |
| S54 | *Paenarthrobacter*<br>*ureafaciens* | 95.21 | 1.17 | 4.41 | 4.41 | 1 | 63.53 | 4785 | 4683 | 18 | 58 | 25 |
| S56 | *Paenarthrobacter* sp. | 96.39 | 0.29 | 4.20 | 4.20 | 1 | 63.87 | 4247 | 4149 | 18 | 57 | 22 |

<sup>a</sup> The genome sequence data were uploaded to the Type (Strain) Genome Server (TYGS) for a whole genome-based taxonomic analysis (Me Kolthoff and Göker, 2019). Genomic relatedness was determined using average nucleotide identity (ANI) values computed with EzBioCloud (Y et al., 2017).

The genomic properties of all of the selected bacterial strains were similar to those of other strains belonging to the same species deposited in GenBank (data not shown).

The TYGS genome-based pipeline was used to refine the identification of the ten bacterial strains based on 16S rRNA gene sequencing (Meier-Kolthoff and Göker, 2019; Yoon et al., 2017) (Table 1; Supplementary Table S2). Strains PSE31B, PFE42 and PFE11 were all assigned to the bacterial species *B. velezensis*. The two *Pseudomonas* strains were identified as *P. salmasensis* strain POE54 and *P. simiae* strain POE78A. Two of the three bacterial strains in the Microccoccaceae family were identified at the species level as *Glutamicibacter halophytocola* strain PFE44 and *Paenarthrobacter ureafaciens* strain S54. However, strain S56 could only be identified at the genus level as *Paenarthrobacter* sp. with a dDDH value of 24% when compared to the closest genome reference *P. ureafaciens* DSM 20126^T^; consequently, this strain may belong to a new species. Similar results were obtained for *Chryseobacterium* sp. POE47 and *Leclercia* sp. strains S52 (*Enterobacter* sp. by 16S rRNA gene sequencing) when compared to the closest phylogenetic species *Chryseobacterium taeanense* DSM 17071^T^ and *Leclercia tamurae* H6S3^T^, for which the corresponding dDDH values were 31.2% and 51.4%, respectively. The calculated ANI values for each strain with the closest related species supported these conclusions (Supplementary Table S2). Table 2 summarizes the general genomic characteristics of the sequenced strains.

Coding sequences were extracted from the genomes of the ten selected strains and classified using the Clusters of Orthologous Groups of proteins (COG) database, revealing four main functional gene classes that were present in all ten genomes (Supplementary Figure S3): (i) Amino acid transport and metabolism, (ii) Carbohydrate transport and metabolism, (iii) Cell wall/membrane/envelope biogenesis, and (iv) Transcription. Other highly represented gene classes included translation, ribosomal structure and biogenesis, signal transduction mechanisms, energy production and conversion, coenzyme transport and metabolism, lipid transport metabolism, and inorganic ion transport and metabolism. No genes in the chromatin structure and dynamics or nuclear structure categories were detected in any genome (Supplementary Figure S3). The “amino acid transport and metabolism” class was the most strongly represent among the COGs in the three members of the *Bacillus* genus, the two members of the *Pseudomonas* genus, and *G. halophytocola* strain PFE44. The most abundant gene families identified in *Chryseobacterium sp.* POE47 were associated with “cell wall/membrane/envelope biogenesis”, while the most strongly represented COG family for *Leclercia sp.* S52, *P. ureafaciens* S54 and *Paenarthrobacter sp.* S56 was “carbohydrate transport and metabolism”. Further details of the COG analysis are presented in Supplementary Figure S3.

In the KEGG analysis, genes associated with “protein families: genetic information processing” were most abundant in the *Bacillus* strains, *Chryseobacterium* sp, POE47 and *Leclercia sp.* S52. For the two *Pseudomonas* strains, the “environmental information processing” category was dominant, while “carbohydrate metabolism” was dominant in the three members of the Micrococcaceae family (Supplementary Figure S4).

At the protein level, analysis with OrthoVenn 3 (https://orthovenn3.bioinfotoolkits.net) revealed that the highest number of orthologous clusters was found in *Pseudomonas* species POE78A and POE54 (3874 and 3826), followed by *B. velezensis* PFE11, PFE42 and PSE31B (3273, 3253 and 3216). The lowest number was recorded in *Chryseobacterium sp.* POE47 (1679) (Supplementary Figure S5). The two pseudomonads shared 1387 clusters, while the three *B. velezensis* strains shared 1377. The two *Paenarthrobacter* strains (S54 and S56) shared 624 clusters, and 483 were also shared with the other strain belonging to the Micrococcaceae family (i.e. *G. halophytocola* PFE44). In total, 530 orthologous clusters were shared by all ten selected strains. A total of 329 gene clusters were specific to a single genome. Of these clusters, 101 belonged to *Leclercia sp.* S52 and 228 were from *Chryseobacterium sp.* POE47 (Supplementary Figure S5).

### 3.5 Genes related to potential PGP and biocontrol traits

A genome annotation analysis conducted using the PGPT-Pred function revealed genomic features associated with plant growth promotion. The genomes of all ten selected strains had similar PGPT classes, with minor differences across the diverse taxa. In all strains, the class with the highest proportion of genes was “Colonizing plant system”, followed by “Stress control and Biocontrol”, “Competitive exclusion”, “Biofertilization”, “Phytohormone and Plant Signaling", “Bioremediation” and “Plant immune response stimulation” (Supplementary Figure S6). A more detailed investigation of PGPT categories highlighted traits of interest linked to direct and indirect effect on plant growth (Figure 3). Supplementary Table S3 shows the full set of genes found in the ten genomes.

**Figure 3.**
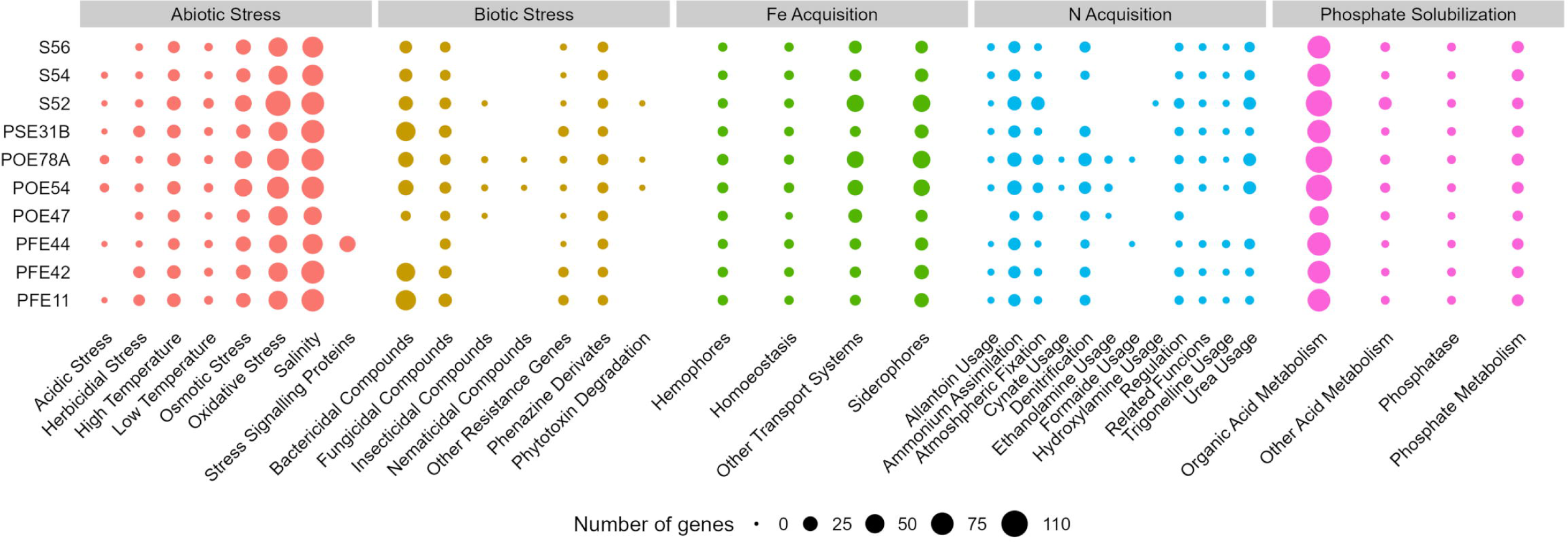
Number of genes of selected PGPT classes related to plant growth promotion and stress control in the genomes of the ten strains. PGPTs were predicted using PGPT-Pred module of PLaBAse v1.01 (Patz et al., 2021). S56, *Paenarthrobacter* sp.; S54, *P. ureafaciens*; S52, *Leclercia* sp.; PSE31B, *Bacillus velezensis*; POE78A, *Pseudomonas simiae*; POE54, *P. salmasensis*; POE47, *Chryseobacterium* sp.; PFE44, *Glutamicibacter halophytocola*; PFE42, *B. velezensis*; PFE11, *B. velezensis*.

Annotation of the genomes against the ’plant bacterial only interaction factors (proteins) (PIFAR)’ dataset revealed two distinct clusters among the bacterial endophytes: one comprising those exhibiting antimicrobial activity and the other comprising those that did not. Minor groups were further segregated based on their taxonomic affiliation (Figure 4). The two pseudomonad strains *P. salmasensis* POE54 and *P. simiae* POE78A belonged to the first group and had the highest percentages of toxin-related factors, which accounted for 44% and 42% of the identified PIFAR, respectively (Figure 4). The three *B. velezensis* strains (PSE31B, PFE42 and PFE11) formed a second distinct cluster with toxin factor percentages ranging from 35 to 38% (Figure 4). *Leclercia* sp. strain S52 clustered more closely with, but separately, to *Bacillus* spp. The lowest toxin content was detected in the *Paenarthrobacter* strains S54 and S56 (24 and 25%), which formed a cluster with *G. halophytocola* PFE44 (Figure 4). These strains belonging to the Micrococcaceae family had the highest content of hormone-related factors (18%). In the remaining strains, hormone-related factors comprised only 7 to 10% of the total PIFAR (Figure 4). EPS was the second most abundant class of factors across all genomes except *Chryseobacterium* sp. POE47, where EPS was the most abundant class, accounting for 29% of the total detected factors (Figure 4). This strain clustered separately from the others. Detoxification related factors comprised a significant proportion of the genomes of all ten strains but were most abundant in *Chryseobacterium sp.* POE47 (14%) and least abundant in the *B. velezensis* strains (7-8%) (Figure 4). Supplementary Table S4 provides detailed information on the predicted PIFAR.

**Figure 4.**
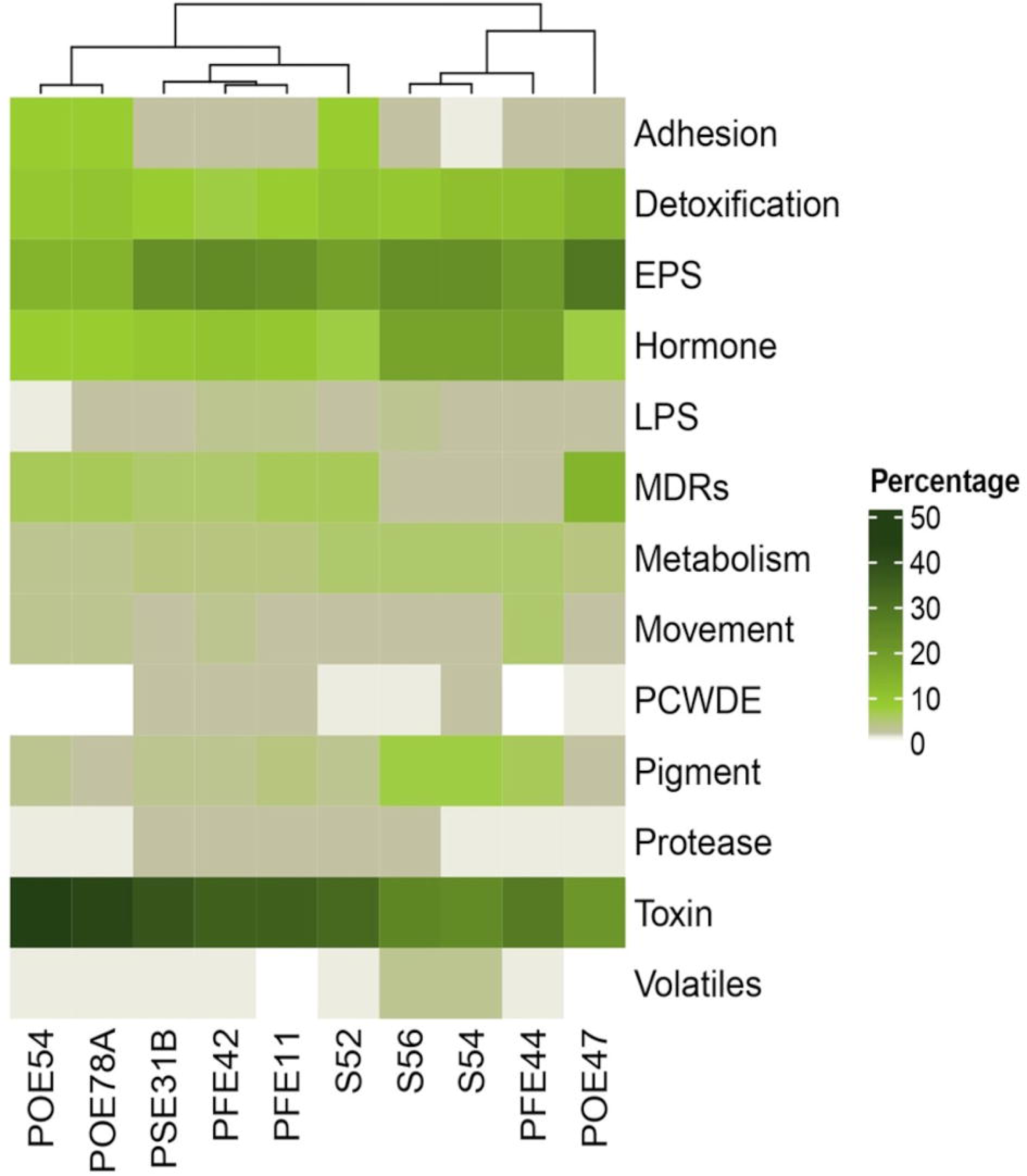
Percentage of genes encoding for ’plant bacterial only interaction factors (proteins)’ predicted through PIFAR-BASE (Patz et al., 2021). Factor categories are specified in the right. Strain ID is indicated in the bottom. POE54, *Pseudomonas salmasensis*; POE78A, *P. simiae*; PSE31B, *Bacillus velezensis*; PFE42, *B. velezensis*; PFE11, *B. velezensis*; S52, *Leclercia* sp.; S56, *Paenarthrobacter* sp.; S54, *P. ureafaciens*; PFE44, *Glutamicibacter halophytocola*; POE47, *Chryseobacterium* sp.

### 3.6 Biosynthetic gene cluster (BGC) mining

AntiSMASH 7.0 database analysis for secondary metabolites (Blin et al., 2023) based on the number of Biosynthetic Gene Clusters (BGCs) in each genome revealed a group comprising *Bacillus* and *Pseudomonas* strains with 11 to 14 BGCs, as shown in Table 3. The three members of the Micrococcaceae family (PFE44, S54 and S56) had six to eight BGCs, while *Chryseobacterium sp.* POE27 and *Leclercia sp.* S52, had the fewest BGCs (three and four, respectively).

**Table 3.** Bacterial Biosynthetic Gene Clusters (BGCs) for secondary metabolites predicted by AntiSMASH 7.0.

| Strain | Predicted BGCs | Total lenght (nt) | % on the total genome |
| --- | --- | --- | --- |
| *Pseudomonas salmasensis* POE54 | 14 | 452,481 | 7.43 |
| *Pseudomonas simiae* POE78A | 13 | 401,988 | 6.46 |
| *Chryseobacterium* sp. POE47 | 3 | 107,090 | 2.43 |
| *Leclercia* sp. S52 | 4 | 147,037 | 2.98 |
| *Bacillus velezensis* PSE31B | 11 | 713,844 | 17.75 |
| *Bacillus velezensis* PFE42 | 12 | 731,931 | 17.39 |
| *Bacillus velezensis* PFE11 | 13 | 761,807 | 19.09 |
| *Glutamicibacter halophytocola* PFE44 | 6 | 149,194 | 4.01 |
| *Paenarthrobacter ureafaciens* S54 | 7 | 175,824 | 3.99 |
| *Paenarthrobacter* sp. S56 | 8 | 183,353 | 4.37 |

The percentage of the genome encoding BGCs was highest in *Bacillus* strains (17-19%), followed by *Pseudomonas* strains (6-7%). The lowest BGC percentages were observed for *Chryseobacterium sp.* POE47 (Table 3). *P.* salmasensis POE54 had the most BGCs (14), with nine types being represented including NRPS (3), NRPS-like (1), RiPP-like (3), arylpolyene (1), betalactone (1), NI-siderophore (1), NAGGN (1), RRE-containing and hybrid (2) (Supplementary Table S5). Moreover, region 13 of this strain’s genome exhibited 100% similarity with the BGC (GenBank: KX931446.1) responsible for obafluorin biosynthesis. No putative metabolites could be identified for the other clusters by MIBiG database comparisons. BGCs associated with surfactin, fengycin, and bacilysin were detected in all *Bacillus* strains. Other identified BGCs were associated with various polyketides including difficidin, bacillaene, and macrolactin, and with the siderophore bacillibactin. *Bacillus* and *Pseudomonas* had higher numbers of PKS and hybrid clusters, and RiPP-like clusters, respectively, than the other strains (Supplementary Table S5). A full list of the antiSMASH results can be found in Supplementary Table S5.

### 3.7 *In vitro* biocontrol activity against bacterial and fungal tomato pathogens

*P. salmasensis* strain POE54 and *P. simiae* strain POE78A, the three *B. velezenzis* strains PSE31B, PFE42 and PFE11 and *Leclercia* sp. strain S52 showed broad antagonistic activity against the three bacterial and two fungal tomato pathogens used in the antimicrobial assays (Table 4; Figures 5 and 6). *Chryseobacterium* sp. strain POE47 only showed antagonistic activity against Forl PVCT127. The three bacterial strains belonging to the genus Micrococcales, i.e. *G. halophytocola* PFE44*, P. ureafaciens* S54 and *Paenarthrobacter* sp. S56, generally showed no antagonistic activity although *Paenarthrobacter sp.* S56 displayed minor activity against Forl PVCT127 (PGI, 21.1%) (Table 4; Figures 5 and 6).

**Figure 5.**
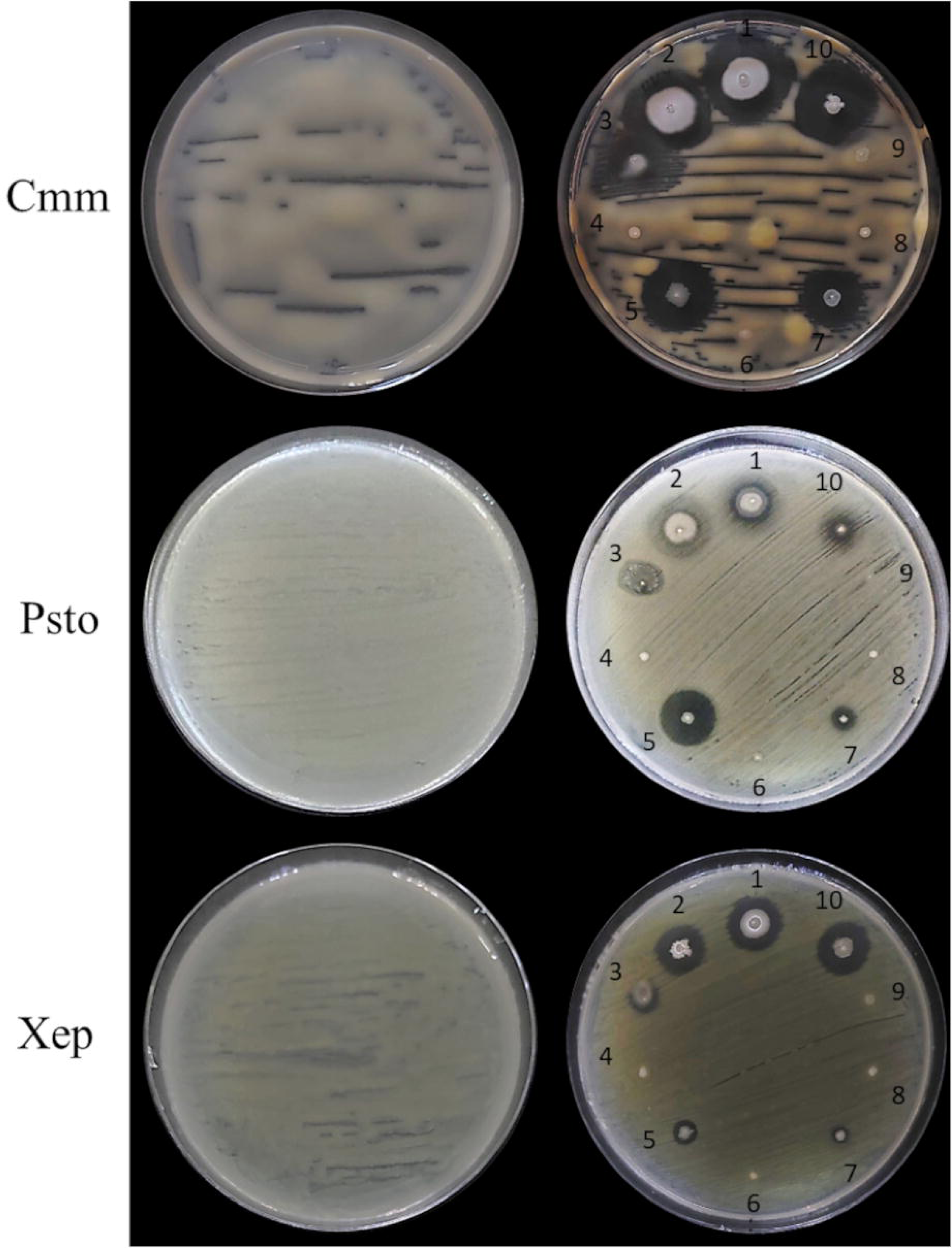
*In vitro* antimicrobial activity of the endophytic bacterial strains against tomato bacterial pathogens. Endophytic bacterial strains: 1, *Bacillus velezensis* PSE31B; 2, *B. velezensis* PFE42; 3, *Leclercia* sp. S52; 4, *Paenarthrobacter ureafaciens* S54; 5, *Pseudomonas salmasensis* POE54; 6, *Chryseobacterium* sp. POE47; 7, *Pseudomonas simiae* POE78A; 8, *Paenarthrobacter* sp. S56; 9, *Glutamicibacter halophytocola* PFE44; 10, *B. velezensis* PFE11. Bacterial pathogens: Cmm, *Clavibacter michiganensis* subsp. *michiganensis* strain PVCT 156.1.1; Psto, *Pseudomonas syringae* pv. *tomato* strain PVCT 28.3.1; Xep, *Xanthomonas euvesicatoria* pv. *perforans* strain NCPPB4321.

**Figure 6.**
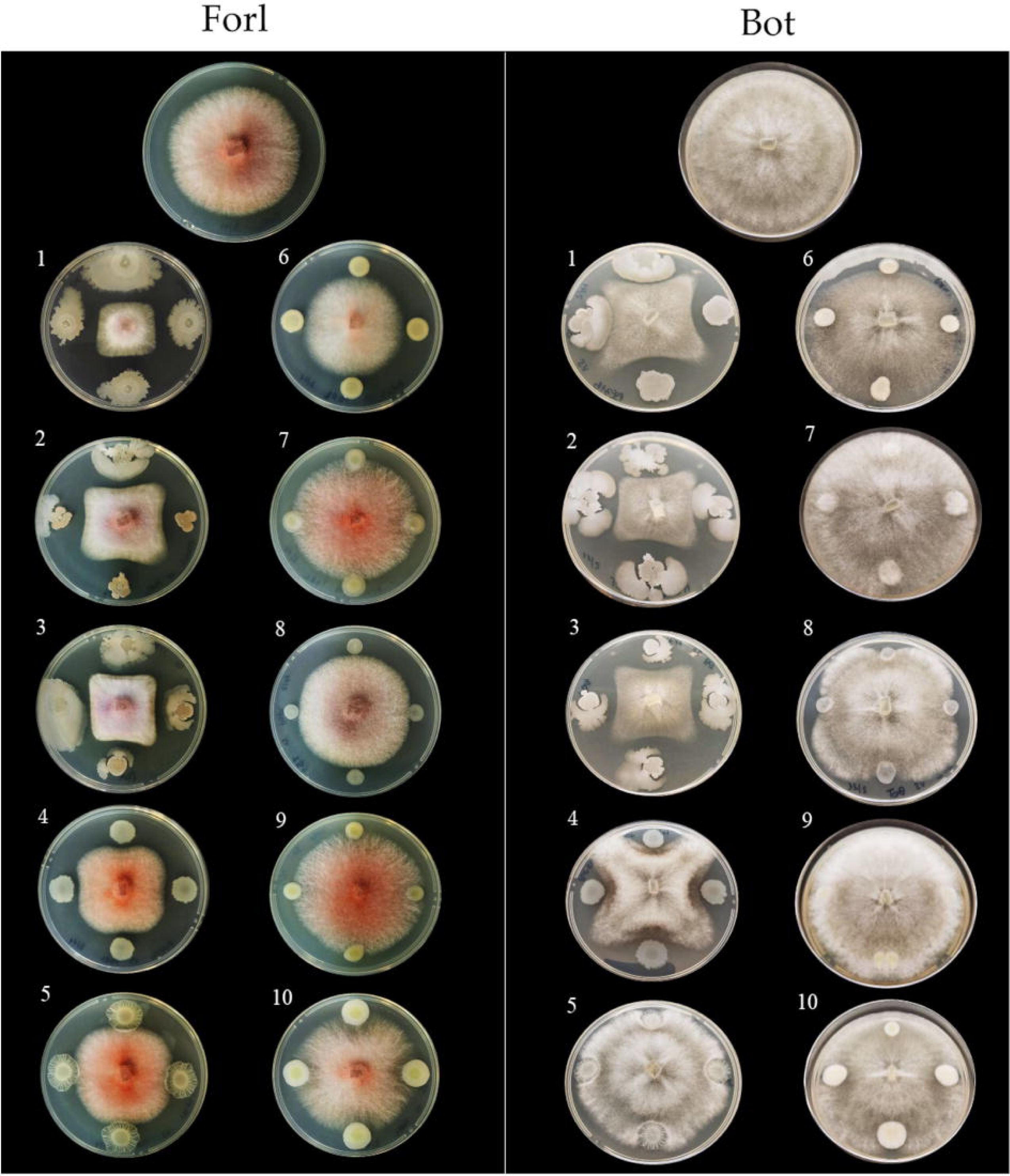
*In vitro* antimicrobial activity of the endophytic bacterial strains against tomato fungal pathogens. Endophytic bacterial strains: 1, *Bacillus velezensis* PSE31B; 2, *B. velezensis* PFE42; 3, *B. velezensis* PFE11; 4, *Pseudomonas salmasensis* POE54; 5, *P. simiae* POE78A; 6, *Chryseobacterium* sp. POE47; 7, *Glutamicicbacter halophytocola* PFE44; 8, *Leclercia* sp. S52; 9, *Paenarthrobacter ureafaciens* S54; 10, *Paenarthrobacter* sp. S56. Fungal pathogens: Forl, *Fusarium oxysporum* f. sp. *radicis-lycopersici* strain PVCT127; Bot, *Botrytis cinerea* strain Bc5.

**Table 4.** *In vitro* antimicrobial activity of the selected bacterial strains against bacterial and fungal tomato pathogens.

| Strain | Inhibition halos<br>(radius, mm) <sup>a</sup> |  |  | Inhibition halos + colony<br>(radius, mm) <sup>a</sup> |  |  | PGI <sup>a,b</sup> |  |
| --- | --- | --- | --- | --- | --- | --- | --- | --- |
|  | Cmm | Psto | Xep | Cmm | Psto | Xep | Forl | Bot |
| *Pseudomonas salmasensis* POE54 | 10.6 ±<br>1.5 | 9.3 ±<br>0.7 | 3.7 ±<br>1.3 | 15.2 ±<br>1.2 | 11.6 ±<br>0.9 | 7.0 ±<br>3.1 | 27.3 ±<br>1.5 | 45.0 ±<br>5.2 |
| *Pseudomonas simiae* POE78A | 8.9 ±<br>1.9 | 3.6 ±<br>0.8 | 1.9 ±<br>1.5 | 12.1 ±<br>1.9 | 4.7 ±<br>1.0 | 3.9 ±<br>1.1 | 23.8 ±<br>4.8 | 18.2 ±<br>10.4 |
| *Chryseobacterium* sp. POE47 | - | - | - | 2.7 ±<br>0.2 | 1.7 ±<br>0.3 | 1.7 ±<br>0.5 | 22.2 ±<br>6.5 | - |
| *Leclercia* sp. S52 | 11.1 ±<br>4.5 | 1.9 ±<br>0.3 | 4.3 ±<br>1.4 | 16.2 ±<br>2.3 | 7.2 ±<br>2.1 | 8.6 ±<br>0.6 | 9.7 ±<br>3.2 | 37.2 ±<br>8.8 |
| *Bacillus velezensis* PSE31B | 10.1 ±<br>0.6 | 2.8 ±<br>0.2 | 5.3 ±<br>1.2 | 18.4 ±<br>0.9 | 7.6 ±<br>1.0 | 12.6 ±<br>3.0 | 59.2 ±<br>2.0 | 57.9 ±<br>4.4 |
| *Bacillus velezensis* PFE42 | 8.7 ±<br>0.8 | 2.8 ±<br>0.4 | 4.3 ±<br>2.3 | 21.6 ±<br>4.1 | 6.2 ±<br>1.3 | 12.9 ±<br>3.1 | 40.6 ±<br>3.0 | 63.6 ±<br>6.6 |
| *Bacillus velezensis* PFE11 | 13.8 ±<br>0.9 | 3.8 ±<br>0.3 | 5.9 ±<br>1.2 | 19.5 ±<br>4.6 | 5.5 ±<br>0.2 | 10.3 ±<br>1.1 | 48.3 ±<br>1.7 | 61.1 ±<br>5.8 |
| *Glutamicibacter halophytocola* PFE44 | - | - | - | 2.6 ±<br>0.5 | 2.0 ±<br>0.7 | 2.6 ±<br>0.3 | - | - |
| *Paenarthrobacter ureafaciens* S54 | - | - | - | 2.8 ±<br>0.2 | 1.9 ±<br>0.4 | 2.5 ±<br>0.4 | - | - |
| *Paenarthrobacter* sp. S56 | - | - | - | 2.4 ±<br>0.4 | 1.6 ±<br>0.3 | 2.1 ±<br>0.3 | 21.1 ±<br>2.8 | - |

Cmm, *Clavibacter michiganensis* subsp. *michiganensis*; Psto, *Pseudomonas syringae* pv. *tomato*; Xep, *Xanthomonas* *euvesicatoria* pv. *perforans*, Forl, *Fusarium oxysporum* f. sp. *radicis-lycopersici*; Bot, *Botrytis cinerea*. <sup>a</sup> Values represent the mean of three replicates. <sup>a</sup> Percentage of Growth Inhibition (Vincent, 1947).

Figures 5 and 6 show the results of the antimicrobial activity assays against bacterial and fungal tomato pathogens. It is particularly notable that the *Bacillus* colonies expanded in the medium, especially when challenged with the fungal pathogens (Figure 6) and the Gram-positive bacterium Cmm strain PVCT 156.1.1 (Figure 5). Based on the radius of the inhibition halo, i.e. the region where microbial growth is absent, *P. salmasensis* strain POE54 showed the greatest inhibitory activity against Psto strain PVCT 28.3.1 (Table 4; Figure 5).

### 3.8 *In planta* biocontrol of bacterial and fungal diseases

Tomato plants treated with each of the ten selected endophytic bacterial strains were challenged with the fungal pathogen *F. oxysporum* f.sp. *radicis lycopersici* strain PVCT127 by inoculation in the soil near the plant at the plant base (i.e. the crown) or with the bacterium *X. euvesicatoria* pv. *perforans* strain NCPPB 4321 by spray inoculation on the epigeal plant portion.

Forty-five days after inoculation, fusarium crown and root rot symptoms were evaluated in longitudinal sections of tomato plants running through the stem to the taproot. All control plants showed dark brown discoloration of the vascular tissues at the stem base extending about 3 cm above soil level and abundant production of adventitious roots (Table 5; Figure 7A). With the exception of the *P. ureafaciens* S54 treatment, all bacterial treatments reduced the percentage of infected plants and significantly reduced disease severity based on the length of discoloration at the crown base (p<0.001) (Table 5). The highest biocontrol efficacy (93.81%) was achieved by using *B. velezensis* PSE31B; only 33% of the plants treated with this strain had vascular discolorations and where discoloration was observed it extended for less than 0.2 cm (meaning that browning was observed only around the inoculation wounds) (Table 5; Figure 7B). Notably, even bacterial strains that showed no antagonistic activity *in vitro*, namely *G. halophytocola* PFE44 and *P. ureafaciens* S54, exerted significant biocontrol over fusarium crown and root rot.

**Figure 7.**
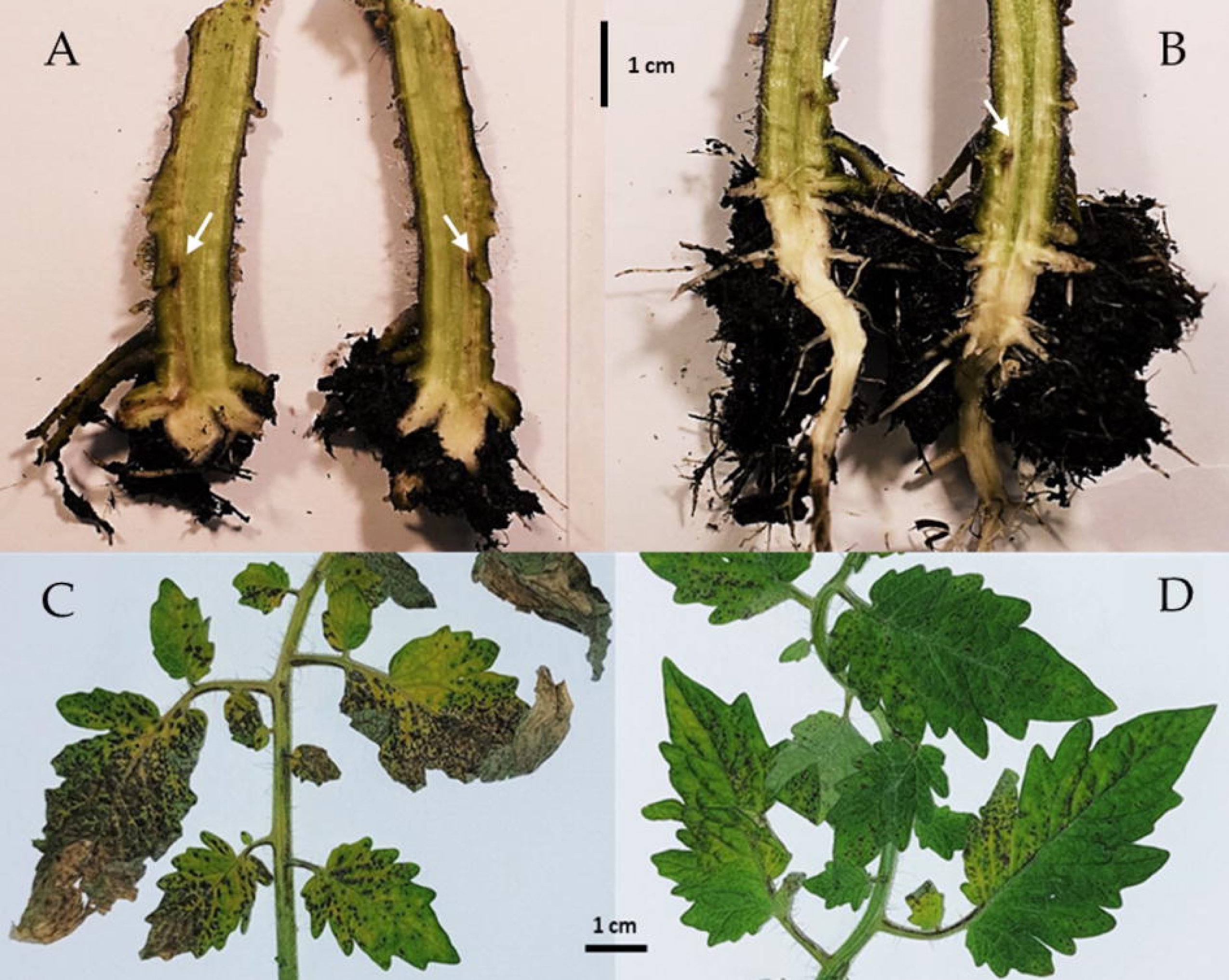
Symptoms of vascular discoloration in tomato plants artificially inoculated with *Fusarium oxysporum* f. sp. *radicis-lycopersici* PVCT127 (A) and in *Bacillus velezensis* PSE31B treated plants (B). Arrows highlight the artificial wound to assist pathogen penetration. Bacterial spot symptoms in tomato plants artificially inoculated with *Xanthomonas euvesicatoria* pv. *perforans* strain NCPPB4321 (C) and in *Pseudomonas simiae* POE78A treated plants (D). In both trials bacterial endophytes were applied by seed soaking and by soil drenching three weeks after plant’s emergence and 72 and 24 h before pathogens inoculation (*Fusarium* and *Xanthomonas*, respectively).

**Table 5.** Biocontrol efficacy of the endophytic bacterial strains against *Fusarium oxysporum* f. sp*. radicis-lycopersici* PVCT127 (Forl) and *Xanthomonas euvesicatoria* pv. *perforans* NCPPB4321 (Xep) in tomato plants in growth chamber. Bacterial strains were applied by seed soaking and by soil drenching three weeks after plant’s emergence and 72 and 24 h before Forl and Xep artificial inoculation, respectively.

| Treatment | Forl |  |  | Xep |  |  |
| --- | --- | --- | --- | --- | --- | --- |
|  | Disease Incidence (%) | Disease severity <sup>a</sup> | Biocontrol efficacy (%) <sup>b</sup> | Disease Incidence (%) | Disease severity <sup>c</sup> | Biocontrol efficacy (%) |
| Control | 100.00 a | 2.73 a | - | 94.68 a | 14.95 a | - |
| *Pseudomonas salmasensis* POE54 | 66.67 a | 0.94 b | 65.44 | 73.06 ab | 3.22 bc | 78.48 |
| *Pseudomonas simiae* POE78A | 83.33 a | 0.86 b | 68.45 | 82.49 ab | 1.55 c | 89.63 |
| *Chryseobacterium* sp. POE47 | 66.67 a | 0.87 b | 68.23 | 74.76 ab | 4.35 bc | 70.91 |
| *Leclercia* sp. S52 | 50.00 a | 0.58 b | 78.78 | 52.94 b | 2.27 c | 84.83 |
| *Bacillus velezensis* PSE31B | 33.33 a | 0.17 b | 93.81 | 81.88 ab | 2.80 c | 81.29 |
| *Bacillus velezensis* PFE42 | 83.33 a | 0.82 b | 69.90 | 84.48 ab | 4.29 bc | 71.27 |
| *Bacillus velezensis* PFE11 | 66.67 a | 0.57 b | 79.17 | 77.20 ab | 2.29 c | 84.66 |
| *Glutamicibacter halophytocola* PFE44 | 66.67 a | 1.04 b | 61.97 | 74.93 ab | 3.94 bc | 73.65 |
| *Paenarthrobacter ureafaciens* S54 | 100.00 a | 0.93 b | 66.06 | 92.41 ab | 11.28 ab | 24.50 |
| *Paenarthrobacter* sp. S56 | 66.67 a | 1.09 b | 60.22 | 81.94 ab | 8.31 abc | 44.44 |
| P value <sup>e</sup> | 0.092 | <0.001 | - | 0.091 | <0.001 | - |

<sup>a</sup> Values represent the length (cm) of the vascular discoloration along the stem. <sup>b</sup> Values express the reduction of the vascular discoloration in plants treated with PGPRs compared to the control. <sup>c</sup> Values represent the percent leaf area affected by bacterial spot. <sup>d</sup> Values express the reduction of area affected by bacterial spot in plants treated with PGPRs compared to the control. <sup>e</sup> Values (mean) in the same column followed by the same letter do not significantly differ based on post-hoc Tukey HSD test at P = 0.05.

Symptoms of bacterial spot observed six days after bacterial inoculation included both pinpoint necrotic spots and/or larger irregular spots that in some cases converged into larger lesions surrounded by chlorotic areas (Figure 7C). Disease incidence (as measured by the percentage of symptomatic leaflets) was not affected by treatment with any endophytic strain except for *Leclercia* sp. S52 (Table 5). However, disease severity (i.e., the percentage of the leaf area containing bacterial spots) was reduced by every bacterial treatment other than those using *P. ureafaciens* strain S54 or *Paenarthrobacter* sp. strain S56 (Table 5). The highest biocontrol efficacy (89.63%) was obtained using *P. simiae* POE78A (Table 5; Fig. 7D).

### 3.9 Growth promotion in tomato nursery plantlets

All of the endophytic bacterial treatments other than those using *Leclercia* sp. strain S52 resulted in growth-promoting activity, although the effects were not always statistically significant. In particular, treatment with *B. velezensis* strains PSE31B, PFE42 and PFE11, *P. salmasensis* POE54 and *Paenarthrobacter sp.* S56 significantly enhanced the height of nursery plantlets at all monitoring time points (p<0.0001) (Figure 8). These bacterial strains also positively influenced the fresh and dry biomasses of roots and shoots (Table 6). The shoot fresh weight was significantly higher in plants treated with *B. velezensis* strains, *P. salmasensis* POE54, and *Paenarthrobacter sp.* S56 (p<0.001), although the treatments did not significantly increase the shoot dry weight in all cases (Table 6). Plants treated with the strains *B. velezensis* PSE31B and PFE11 and *Paenarthrobacter sp.* S56 also had significantly increased fresh and/or dry root weights (p=0.001) (Table 6).

**Figure 8.**
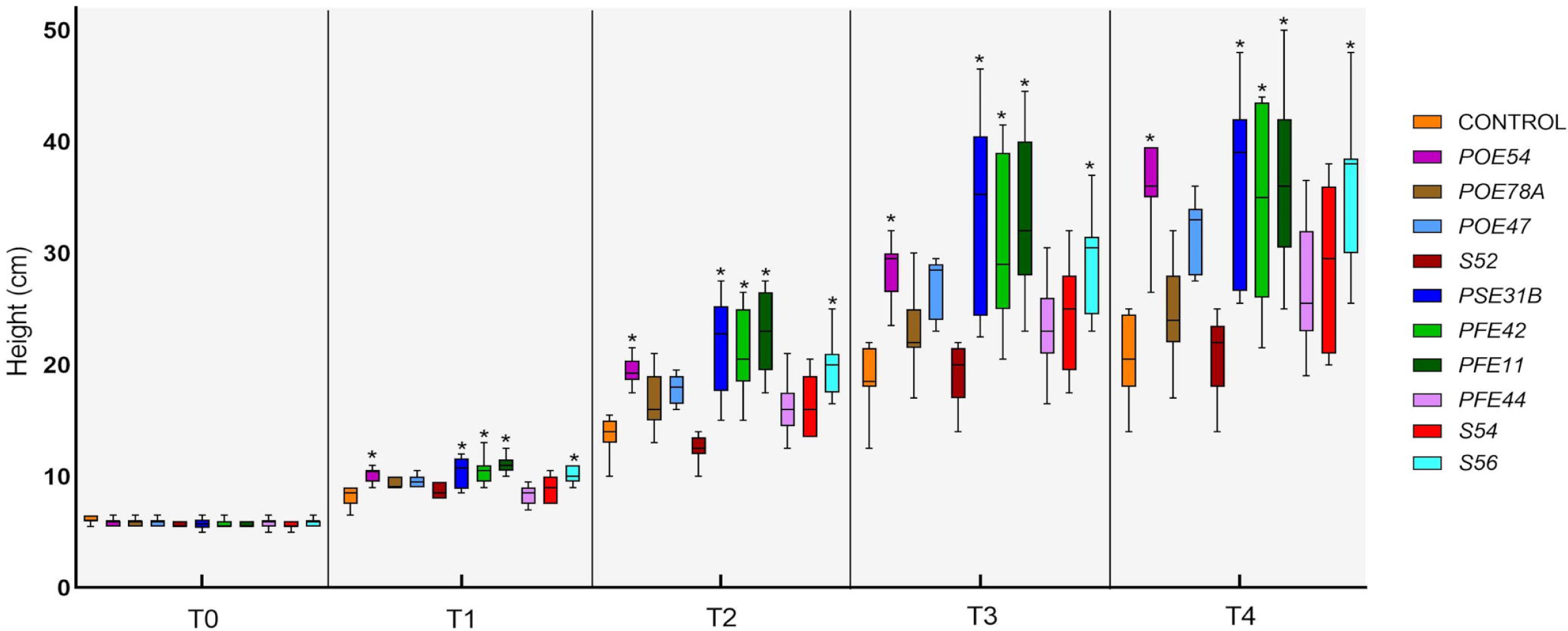
Time-course evaluation of tomato plant height in the PGP trial in growth chamber: T0 (before the treatments), T1-4 (1-4 weeks after the treatments). Plants were treated by soil drenching immediately after transplant. Asterisks denote statistical significance compared to the not treated plants (Control) based on post-hoc Tukey HSD test at P = 0.05. POE54, *Pseudomonas salmasensis*; POE78A, *P. simiae*; POE47, *Chryseobacterium* sp.; S52, *Leclercia* sp.; PSE31B, *Bacillus velezensis*; PFE42, *B. velezensis*; PFE11, *B. velezensis*; PFE44, *Glutamicibacter halophytocola*; S54, *Paenarthrobacter ureafaciens*; S56, *Paenarthrobacter* sp.

**Table 6.**
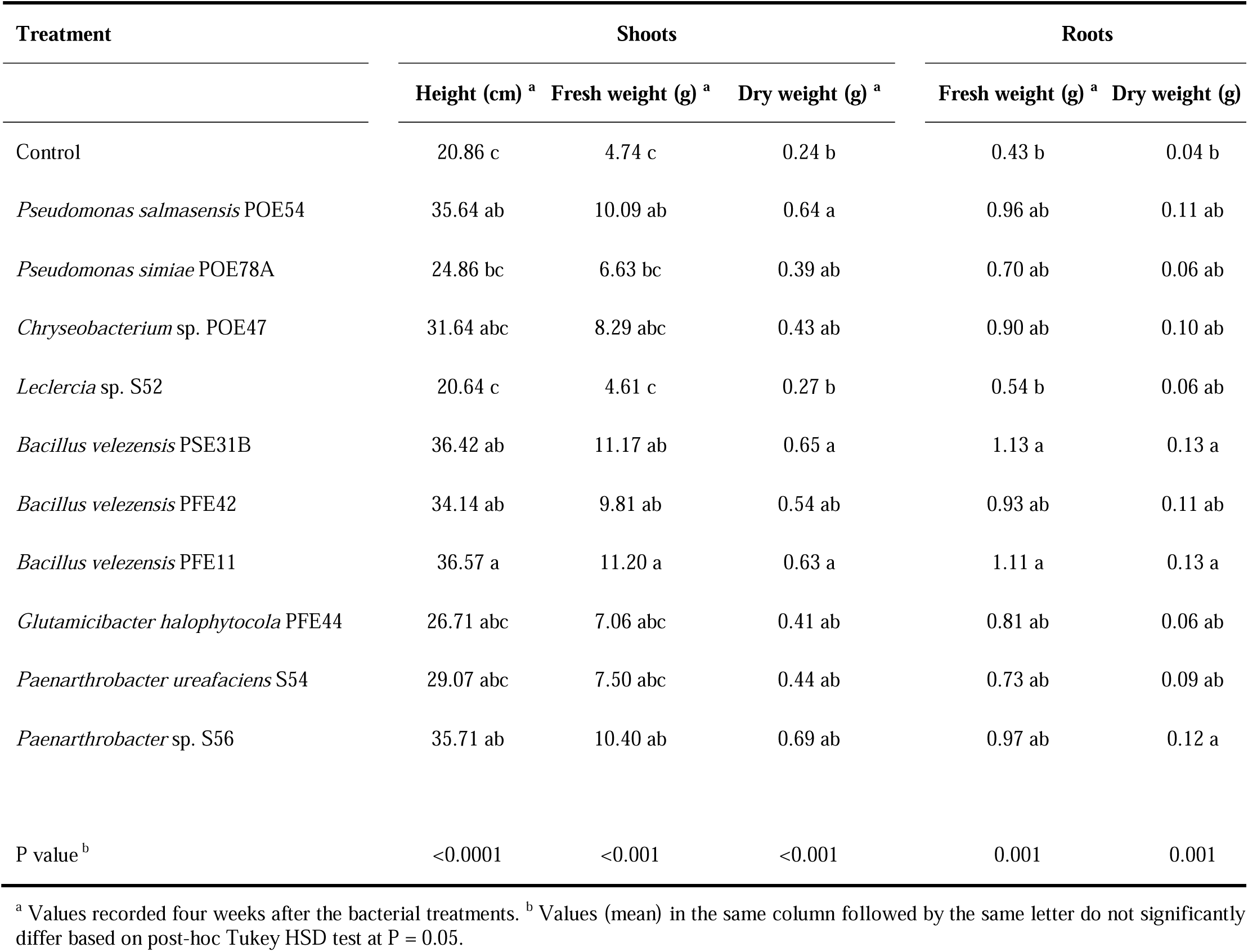
Plant growth promotion efficacy of the endophytic bacterial strains in tomato plants in growth chamber. Bacterial strains were applied by soil drenching after transplant.

| Treatment | Shoots |  |  | Roots |  |
| --- | --- | --- | --- | --- | --- |
|  | Height (cm) <sup>a</sup> | Fresh weight (g) <sup>a</sup> | Dry weight (g) <sup>a</sup> | Fresh weight (g) <sup>a</sup> | Dry weight (g) |
| Control | 20.86 c | 4.74 c | 0.24 b | 0.43 b | 0.04 b |
| *Pseudomonas salmasensis* POE54 | 35.64 ab | 10.09 ab | 0.64 a | 0.96 ab | 0.11 ab |
| *Pseudomonas simiae* POE78A | 24.86 bc | 6.63 bc | 0.39 ab | 0.70 ab | 0.06 ab |
| *Chryseobacterium* sp. POE47 | 31.64 abc | 8.29 abc | 0.43 ab | 0.90 ab | 0.10 ab |
| *Leclercia* sp. S52 | 20.64 c | 4.61 c | 0.27 b | 0.54 b | 0.06 ab |
| *Bacillus velezensis* PSE31B | 36.42 ab | 11.17 ab | 0.65 a | 1.13 a | 0.13 a |
| *Bacillus velezensis* PFE42 | 34.14 ab | 9.81 ab | 0.54 ab | 0.93 ab | 0.11 ab |
| *Bacillus velezensis* PFE11 | 36.57 a | 11.20 a | 0.63 a | 1.11 a | 0.13 a |
| *Glutamicibacter halophytocola* PFE44 | 26.71 abc | 7.06 abc | 0.41 ab | 0.81 ab | 0.06 ab |
| *Paenarthrobacter ureafaciens* S54 | 29.07 abc | 7.50 abc | 0.44 ab | 0.73 ab | 0.09 ab |
| *Paenarthrobacter* sp. S56 | 35.71 ab | 10.40 ab | 0.69 ab | 0.97 ab | 0.12 a |
| P value <sup>b</sup> | <0.0001 | <0.001 | <0.001 | 0.001 | 0.001 |

<sup>a</sup> Values recorded four weeks after the bacterial treatments. <sup>b</sup> Values (mean) in the same column followed by the same letter do not significantly differ based on post-hoc Tukey HSD test at P = 0.05.

## 4 Discussion

To identify potential bacterial bioinoculants, a microbiome-guided top-down approach was used to select ten bacterial strains belonging to different taxa from the core microbiome of tomato plants at different stages in the production chain. Bacterial endophytes isolated from tomato seeds and roots were thus not selected on the basis of *in vitro* analyses of phenotypic characteristics associated with PGP activity or antagonism against phytopathogenic microorganisms. This approach resulted in the selection of taxa from the comparatively understudied genera *Leclercia, Chryseobacterium, Glutamicibacter* and *Paenarthorbacter* alongside taxa more commonly used as biofertilizers and biocontrol agents such as *Pseudomonas* and *Bacillus* species. Complete genome sequencing of the strains led to the revision of identifications based on 16S rRNA gene sequences in some cases and made it possible to dissect the strains’ genetic makeup, focusing particularly on phyto-beneficial traits. The resulting information provides valuable insights into the potential biotechnological applications of the strains as efficient bioinoculants for tomato growth and protection in different stages of production.

Microbiome-guided approaches for selecting bacteria beneficial to tomato growth have been used in several studies, only a few of which have focused on the core microbiome (Tian et al., 2017; Bergna et al., 2018). The plant core microbiome is defined as any set of microbial taxa, along with their genomic and functional attributes, that is distinctive to a specific host or environment (Lundberg et al., 2012; Neu et al., 2021). Core microbes create cores of interactions that can be used to optimize microbial functions at the individual plant and ecosystem levels (Toju et al., 2018). Generally, the association between the bacteria in the core microbiome and the potential PGPRs is preceded by a preliminary *in vitro* screening to select strains with the greatest potential as bioinoculants (Tian et al., 2017; Bergna et al., 2018). However, such bottom-up approaches may fail to detect bacteria that do not express the desired characteristics *in vitro* but are nevertheless important representatives of the core microbiome in nature (Compant et al., 2019).

To avoid this problem and select a set of bacteria that is adaptable to diverse conditions, an alternative approach was adopted in this work: instead of performing functional screening first, we began by identifying the core endophytic microbiome of a set of samples representing tomato plants at multiple stages of the production process (Anzalone et al., 2022). Anzalone et al. (2022) had previously studied the composition and shaping of the microbial community in the rhizosphere in tomato plants cultivated with and without soil from nursery growth to the greenhouse, revealing that these bacterial communities were mainly shaped by the substrate or soil in which the plants were grown and differed significantly between plant compartments (rhizosphere and endorhizosphere) and plant growth stages. The most represented taxa were *Rhodococcus* in seeds, *Flavobacterium* in the rhizosphere and *Enterobacter, Pseudomonas* and *Bacillus* in the endorhizospheres of plants grown in peat, agricultural soil and coconut fiber, respectively. A subsequent core microbiome analysis identified the same taxa as the five most abundant ones in the core of the plant-associated samples.

Following a metagenomic analysis of the endosphere of tomato plant seeds and roots, we set up a collection of 94 bacterial endophytes. As ubiquitous colonizers of plants, endophytes are known to strongly influence plant health and productivity; they can have a variety of desirable effects including enhanced plant growth, reduced susceptibility to diseases, and increased tolerance to environmental stresses (Compant et al., 2005; Hardoim et al., 2008; Bulgarelli et al., 2013; Santoyo et al., 2016).

A high proportion of the collected strains showed tolerance to salt stress *in vitro*. This relatively high abundance of salt-tolerant bacterial strains suggests that the typically high salinity of the water and soil in the sampling area enriched the fraction of root endophytes with such tolerance, as previously demonstrated by Flemer et al. (2022). Other beneficial features observed in the collection included siderophore production and solubilization of insoluble organic phosphate; in addition, a few strains were able to produce hydrogen cyanide and ACC deaminase. Moreover, around 30% of the strains showed broad antagonistic activity against five tomato pathogens. Partial 16S rRNA gene sequencing made it possible to provisionally assign most of the strains of the collection to *Bacillus* and *Pseudomonas* spp. These genera are the most highly represented among cultivable endophytic bacteria isolated from different plant species including tomato (Tian et al., 2017; Bergna et al., 2018; Anzalone et al., 2021; Cochard et al., 2022) and from different plant compartments, as reviewed by Riva et al. (2022). Bacteria belonging to the Gram-positive genera *Arthrobacter, Curtobacterium, Glutamicibacter, Microbacterium, Paenarthrobacter, Paenibacillus, Priestia* and *Staphylococcus* and to the Gram-negative genera *Chryseobacterium, Delftia, Enterobacter, Ewingella, Flavobacterium, Serratia* and *Stenotrophomonas* were also part of our collection. Some of these genera have been isolated in other studies on beneficial bacteria (Abbamondi et al., 2016; Tian et al., 2017; Anzalone et al., 2021; Cochard et al., 2022).

Instead of limiting our search for potential beneficial properties to strains well-known for high antagonistic activity *in vitro*, i.e. *Bacillus* and *Pseudomonas* (Tian et al., 2017; Anzalone et al., 2021), we broadened our selection to include less frequently studied genera belonging to the core microbiome. After constructing high-quality genomes for ten strains from the core microbiome that had been assigned to diverse taxa based on preliminary 16S rRNA gene sequencing, we obtained more conclusive species- or genus-level identifications. Specifically, we discovered that the ten selected core microbiome strains belonged to the following taxa: *B. velezensis* (strains PSE31B, PFE42, PFE11), *P. salmasensis* POE54 and *P. simiae* POE78A. Strains *Enterobacter* sp. S52 and *Arthrobacter* sp. S54 were reassigned to *Leclercia* sp. and *P. ureafaciens*, respectively. The taxonomic affiliations of *G. halophytocola* PFE44, *Paenarthrobacter* sp. S56, and *Chryseobacterium* sp. POE47 were confirmed. The last wo strains could only be classified at the genus level by TYGS according to dDDH and ANI values below 70% and 95%, respectively, suggesting that they might represent new bacterial species (Goris et al., 2007; Auch et al., 2010; Meier-Kolthoff et al., 2013)

The ten bacterial strains from the tomato endosphere, although to a different extent, exhibited biocontrol and/or PGP properties, protecting tomato plants against biotic stress and stimulating the growth of tomato seedlings regardless of their activity *in vitro*. These results highlight the limits of *in vitro* screening methods, which often fail to capture the complex interactions that occur in the rhizosphere (Bulgarelli et al., 2013; Mendes et al., 2013). Nutrient availability, soil pH, microbial competition, and plant exudates all influence the efficacy of Plant Growth-Promoting Bacteria (PGPB). Therefore, *in vitro* assays may not accurately predict how bacteria will perform under field conditions (Berg and Smalla, 2009). Furthermore, our strains underwent testing in two distinct pathosystems featuring both belowground and aboveground pathogens. In the trials, the endophytic bacteria were applied to tomato plants by soil drenching and to seeds by seed coating. Importantly, given the diverse biocontrol mechanisms employed by bacteria, which include competing for resources, niche exclusion, and the production of antimicrobial compounds, it is possible that our bacteria may have the capability to induce systemic resistance in the tomato plants (Compant et al., 2005; Lugtenberg and Kamilova, 2009; Singh et al., 2011; Pieterse et al., 2014; Hardoim et al., 2015; Santoyo et al., 2016; Hanifah et al., 2023; Kelbessa et al., 2023).

The selected *Bacillus* and *Pseudomonas* strains, and *Leclercia* sp. strain S52, exhibited broad-spectrum antimicrobial activity against five tomato pathogens *in vitro*. The *Bacillus* endophytes produced the largest inhibition halos when tested against bacterial pathogens and imposed the most significant radial growth inhibition on phytopathogenic fungi. In accordance with this result, five different *Bacillus* species (*B. subtilis, B. velezensis, B. amyloliquefaciens, B. pumilus, B. brevis* and *B. cereus*) were previously shown to exhibit strong antagonism towards diverse phytopathogens (Dimkić et al., 2022; Etesami et al., 2023). This fact together with their high growth rates and tolerance of unfavourable environmental conditions has made them popular biocontrol agents (Fischer et al., 2013). The *B. velezensis* strains (and especially PSE31B) also demonstrated effective control over both tomato fusarium crown and root rot and bacterial spot *in planta*, reaffirming previous reports of this species’ significant biocontrol potential (Felipe et al., 2021; Chen et al., 2022). Notably, *B. velezensis* belongs to the *B. subtilis* group, which contains strains known for their ability to form beneficial associations with plant roots and exert beneficial effects including plant growth promotion and biocontrol of pathogens in several economically important crops, including tomato (Balderas-Ruíz et al., 2021; Felipe et al., 2021; Chen et al., 2022; Mosela et al., 2022). These strains also substantially promoted tomato plant growth in the growth chamber, leading to significantly increased shoot height and fresh and dry weight. These results are consistent with previous reports by Dhouib et al. (2019) and Balderas-Ruíz et al. (2021).

Our genomic findings for the *Bacillus* endophytes are consistent with previous results and align well with the characteristic traits of these species. Their antimicrobial activity may be partly due to their production of diverse bioactive secondary metabolites, which is driven by genes that are located within large genomic islands (BGCs) encoding mega-enzymes including non-ribosomal peptide synthetases and polyketide synthases (Ongena et al., 2007; Rabbee et al., 2019). These metabolites include surfactin and fengycin lipopeptides, which are known to exert antimicrobial activity against various fungal and bacterial pathogens, and to induce systemic resistance and promote biofilm formation (Ongena et al., 2007; Zhao et al., 2017; Penha et al., 2020). A BGC encoding enzymes producing the siderophore bacillibactin, which is highly conserved in the *B. subtilis* group, was also identified (Miethke et al., 2006). This siderophore enables efficient acquisition of Fe^3+^ and other metals in iron-deficient environments, depriving plant pathogens of essential elements (Niehus et al., 2017). Another significant group of BGCs encoded enzymes producing polyketides (e.g. difficidin, bacillaene, and macrolactin) that also play a role in antimicrobial activity (Caulier et al., 2019), and the cluster for the synthesis of bacilysin, a common broad-spectrum antimicrobial dipeptide (Nannan et al., 2021).

Both *P. salmasensis* POE54 and *P. simiae* POE78A displayed effective biocontrol of tomato diseases as well as plant growth promoting activity, with *P. simiae* POE78A exerting the most effective biocontrol activity against *Xanthomonas euvesicatoria* pv. *perforans* in this study. Both *P. salmasensis* and *P. simiae* were identified in a refined taxonomy of the larger *P. fluorescens* complex (Vela et al., 2006; Girard et al., 2021). *Pseudomonas* spp. have multiple traits that make them valuable biocontrol agents, including rapid *in vitro* growth, effective utilization of root exudates, and robust colonization and proliferation in the rhizosphere as well as the production of diverse bioactive metabolites including antibiotics, siderophores, volatiles, and extracellular enzymes, enhancing their competitive edge against other microorganisms (Haas and Défago, 2005; Mercado-Blanco and Bakker, 2007; Loper et al., 2012; Raaijmakers and Mazzola, 2012; Fischer et al., 2013; Höfte, 2021). Many root-associated pseudomonads possess BGCs for the production of antimicrobial compounds such as cyclic lipopeptide biosurfactants, 2,4-diacetylphloroglucinol, phenazines, hydrogen cyanide, and pyrrolnitrin (Loper et al., 2012). *Pseudomonas* strains have also been shown to effectively control *Fusarium oxysporum* f. sp. *radicis-lycopersici* and *Xanthomonas euvesicatoria* pv. *perforans* (Zhang et al., 2015; Felipe et al., 2021; Anzalone et al., 2021).

At least two widely studied *P. fluorescens* biocontrol strains have been reclassified as *P. simiae*, namely WCS417 (Berendsen et al., 2015; Pieterse et al., 2021) and PICF7 (Martínez-García et al., 2015). The phenotypic and genomic determinants of these strains were dissected and they were shown to exhibit biocontrol activity against fungal, bacterial pathogens and nematodes in a wide range of plant species, and to induce systemic resistance in the host and improve tolerance against abiotic stresses (Pieterse et al., 2021). The BGC for obafluorin, a broad-spectrum antibiotic commonly associated with P*. fluorescens*, was found in the genome of POE54 (Wells et al., 1984; Scott et al., 2017). In addition to BGCs responsible for siderophores, the genome of this strain contained many clusters that display no apparent similarity to known BGCs in the MIBiG database and thus warrant further investigation.

*Leclercia* sp. strain S52, which was isolated from tomato seeds, showed good *in vitro* antimicrobial activity, and also efficiently reduced the damage caused by *Fusarium oxysporum* f. sp. *radicis-lycopersici* and *Xanthomonas euvesicatoria* pv. *perforans* during *in vivo* trials but did not positively influence plant growth. Although new rhizosphere-associated *Leclercia* species and related genera have recently been described (Maddock et al., 2022), we were unable to assign our strain to any known species, suggesting that it may represent a new taxonomic entity. Despite ongoing concerns about the use of Enterobacteriaceae species as PGPRs given the existence of human pathogens within this taxon, several studies have investigated this topic (Berg et al., 2005; Erlacher et al., 2015). Many studies have confirmed that Enterobacteriaceae, including *L. adecarboxylata*, are indigenous components of the plant microbiome in different species (Kelemu et al., 2011; Erlacher et al., 2014, 2015; Verma et al., 2015; Shahzad et al., 2017; Tian et al., 2017; Anzalone et al., 2021). In particular, Anzalone et al. (2021) found that 40% of the endophytic bacteria isolated from the tomato endorhizosphere in four tomato farms within the area from which our samples were taken belonged to the Enterobacteriaceae family. *L. adecarboxylata* has previously been shown to promote plant growth (Sarma et al., 2004; Shahzad et al., 2017) and to mitigate both abiotic (Kang et al., 2019, 2021; Ahmed et al., 2021) and biotic (Lee et al., 2023) stresses. AntiSMASH analysis revealed that the genome of the selected *Leclercia* strain contained a BGC for the production of enterobactin, a well-known siderophore with an extraordinary iron affinity (Raymond et al., 2003).

Among the remaining selected strains, *Chryseobacterium* sp. POE47 and *Paenarthrobacter* sp. S56 exhibited limited *in vitro* antagonistic activity against the phytopathogenic fungus *Fusarium oxysporum* f. sp. *radicis-lycopersici* PVCT127 but had no detectable antagonistic effects against the tested phytopathogenic bacteria. Moreover, *G. halophytocola* PFE44 and *P. ureafaciens* S54 exhibited no *in vitro* antagonistic activity at all. In addition, *Chryseobacterium* sp. strain POE47 did not promote tomato plant growth under our experimental conditions. However, in contrast to its poor antimicrobial activity *in vitro* it did effectively control fusarium crown and root rot and bacterial spot *in planta*. In previous studies, *Chryseobacterium* species, formerly classified under the genus *Flavobacterium*, were found to enhance plant growth and exhibit biocontrol activity (Ramos Solano et al., 2008; Sang et al., 2018). *Flavobacterium* strains directly contribute to plant growth by promoting nutrient cycling and supplying beneficial plant hormones (Dardanelli et al., 2010; Sang et al., 2018). A recent study has also explored their halotolerance, suggesting that they might be promising PGPR in saline environments (Jung et al., 2023). Genome mining revealed that these species have a high content of EPS-coding genes, which are known to enhance plant growth and drought tolerance (Naseem et al., 2018). Beneficial traits related to stress relief, biocontrol, biofertilization, and phytohormone production were also identified, in line with the findings of Jung et al. (2023). Interestingly, our genomic analysis showed that this strain had fewer BGCs than any of the other selected strains, with none of them encoding genes producing antimicrobial compounds. This likely explains its poor *in vitro* antagonism.

The strains examined in this work included three bacterial strains belonging to the family Micrococcaceae, a taxon that was reclassified in 2016 (Busse, 2016). This caused its genera to be renamed to *Glutamicibacter, Paeniglutamicibacter, Pseudoglutamicibacter, Paenarthrobacter*, and *Pseudarthrobacter* (Busse, 2016). While both *Glutamicibacter* PFE44 and the two *Paenarthrobacter* strains (S54 and S56) effectively reduced the severity of fusarium crown and root rot symptoms *in planta*, only the *Glutamicibacter* endophyte significantly reduced bacterial spot severity. Both *Paenarthrobacter* and *Glutamicibacter* species have recently emerged as promising PGPR – for example, strains of *P. nitroguajacolicus* mitigated water scarcity in tomato and increased the vigor index values of tomato seedlings (Xiong et al., 2019; Christakis et al., 2021; Fu et al., 2021; Riva et al., 2021; Vasseur-Coronado et al., 2021). Genomic analysis of the Micrococcaceae strains revealed a predominance of genes encoding toxins, exopolysaccharides (EPS), phytohormones, and detoxification products. Moreover, AntiSMASH analysis revealed a BGC encoding desferrioxamine E, a siderophore widely produced by *Streptomyces* and related bacteria (Horinouchi et al., 2010). Desferrioxamine E plays a crucial role in ferric transportation and indirectly acts against fungi (Horinouchi et al., 2010). Additionally, EPS produced by *G. halophytocola* have been shown to mitigate abiotic stresses and promote plant growth (Xiong et al., 2019; Chen et al., 2023).

The overall PGP potential of the ten bacterial strains was evaluated *in silico* by using the PGPT-Pred function of PLa-BAse (Patz et al., 2021) to analyze the prevalence of genes predicting plant growth promoting traits. This tool helped to unveil functional processes responsible for a strain’s PGP activity by revealing groups of genes with direct and indirect beneficial effects on plants. A comprehensive inventory of genes related to colonization, biofertilization, phytohormones, and plant signaling, among other processes was thus obtained for each strain. As noted above, these genes included BGCs encoding antimicrobial substances, chemotaxis- and surface attachment-related genes pivotal for recruitment and colonization in the rhizosphere (Knights et al., 2021), and bacterial secretion systems that play essential roles in out-competing other rhizobacterial strains during root colonization in the host plant (Lugtenberg and Kamilova, 2009; Pieterse et al., 2014; Lucke et al., 2020). The identified genes with direct beneficial effects on plant growth included those influencing traits related to fertilization such as potassium and phosphate solubilization, nitrogen and iron acquisition, sulfur assimilation, and carbon dioxide fixation, all of which enhance nutrient availability (Vessey, 2003; Lugtenberg and Kamilova, 2009; Singh et al., 2011; Das et al., 2022). The strains also carried several functional genes putatively involved in abiotic stress alleviation by stimulating plant immune responses and managing biotic stress via fungicidal and bactericidal activities, indirect traits that foster plant health and growth (Compant et al., 2005; Kelbessa et al., 2023).

The plant interaction traits of the ten selected strains were annotated using PIFAR to support the development of plant bioinoculants that can rapidly establish suitable environments and form beneficial interactions upon application to plants. This analysis clearly separated the strains showing *in vitro* antimicrobial activity from those without such activity. Such differences between bacterial strains could be further exploited to develop bioinoculants with specific target activities, such as biofertilization, biostimulation, or biocontrol.

In conclusion, the studied *Bacillus* and *Pseudomonas* strains exhibited high efficacy both in promoting plant growth and in protecting against pathogens, justifying their predominance in the bioinoculant market (Backer et al., 2018). However, other strains belonging to less known and explored genera also demonstrated effective PGP and BCA activity. The ten bacterial endophytes examined in this work were selected based on an analysis of the core microbiome of tomato seeds and the rhizosphere and endorhizosphere of tomato plants at different stages within the growing chain. Importantly, the collected samples represent a wide range of management practices, environmental conditions, nursery materials (seeds and substrate), and conditions after transplantation into either agricultural soil that had been used for tomato cultivation for several years or soilless crop cultivation media. The genomic data obtained in this work will in future be employed to plan the use of the selected strains individually or in consortia by coupling strains with different traits, effects, and mechanisms of action to obtain synergistic beneficial effects. These strains could potentially be applied as seed dressings or soil drenches at different stages of plant growth, although further research will be needed to develop reliable standardized treatment protocols with predictable effects under specific conditions.

## 6 Author Contributions

Conceptualization and designing the experiment: VC, RRV, DN, FG, AA. Methodology: VC, RRV, DN, FG, SG, AA. Investigation: DN, FG, SG, AA, GD, AM, MEM. Resources: VC, RRV. Writing - original draft preparation: DN. Writing - review and editing: DN, FG, SG, AA, GD, AM, MEM, VC, RRV. Data curation and Formal Analysis: DN, SG, GD, AM. Supervision and project administration: VC, RRV, FG. Funding acquisition: VC, RRV. All authors contributed to the article and approved the submitted version.

## 8 Supplementary Material

The Supplementary Material for this article can be found online at:

## 9 Data Availability Statement

The datasets presented in this study can be found in online repositories. Sequences of the strains used in the present study were submitted to the GenBank database. GenBank accession numbers (MZ066824-MZ066917) are shown in Supplementary Table 1. All the assembled genomes and respective raw reads are available under BioProject Id: PRJNA1096641.

