## Supplementary figures S1-S6 for "Genomic Insights and Biocontrol Potential of Ten Bacterial Strains from the Tomato Core Microbiome"

Supplementary Material

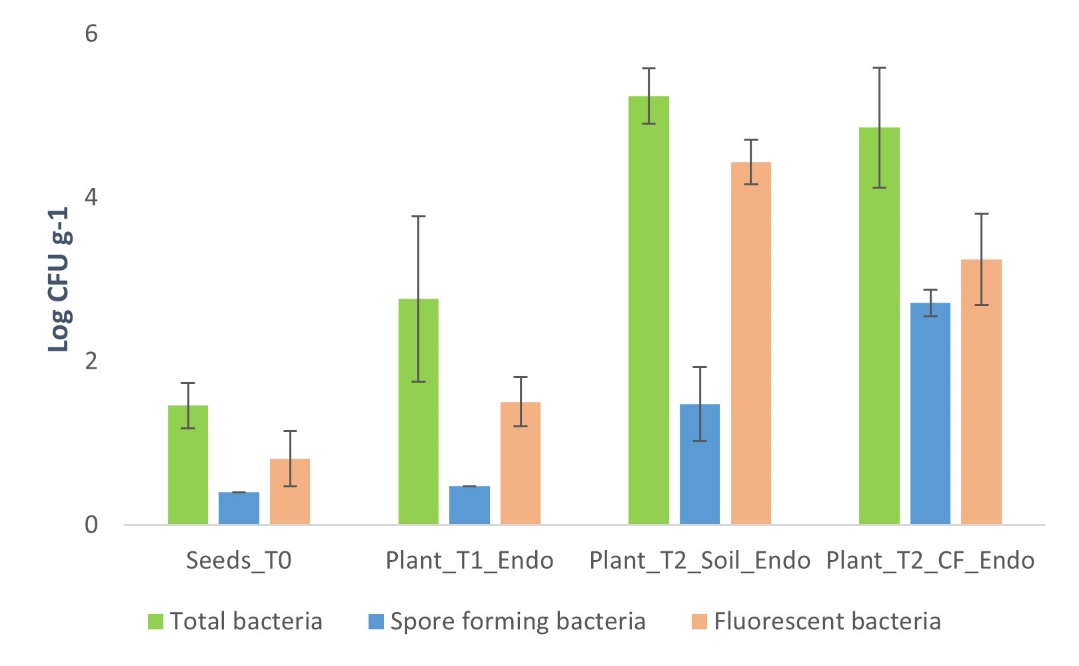

**Supplementary Figure 1.** Total, fluorescent, and spore forming bacterial population size obtained in culture from the different samples: seeds (Seeds_T0), and endorhizosphere of plantlets in the nursery (Plant_T1_Endo) and endorhizosphere of tomato plants in the two cultivation systems, agricultural soil (Plant_T2_Soil_Endo) and coconut fiber substrate (Plant_T2_CF_Endo).

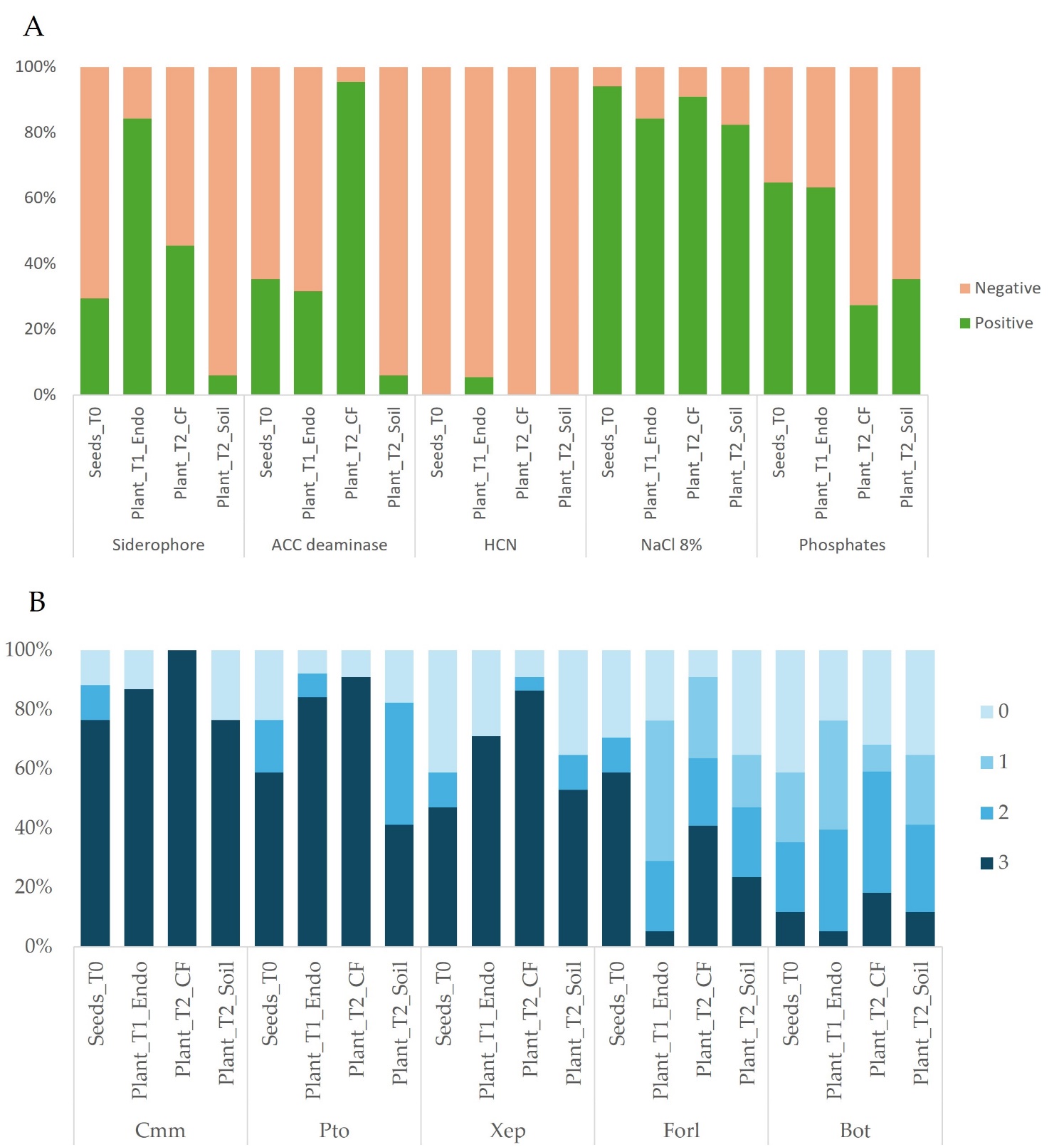

**Supplementary Figure 2.** Characterization of the bacterial strains from the endosphere samples for the (A) presence/absence of PGP traits (B) classes of biocontrol activity. Bacterial inhibition area: 0, no antagonism; 1, small area around the bacterial growth (1-3 mm); 2, large inhibition area (3-10 mm); 3, inhibition of pathogens growth. Fungal Percentage of Growth Inhibition (PGI): 0, no antagonism; 1, PGI <30%; 2, PGI > 30 <60%; 3, PGI>60%.

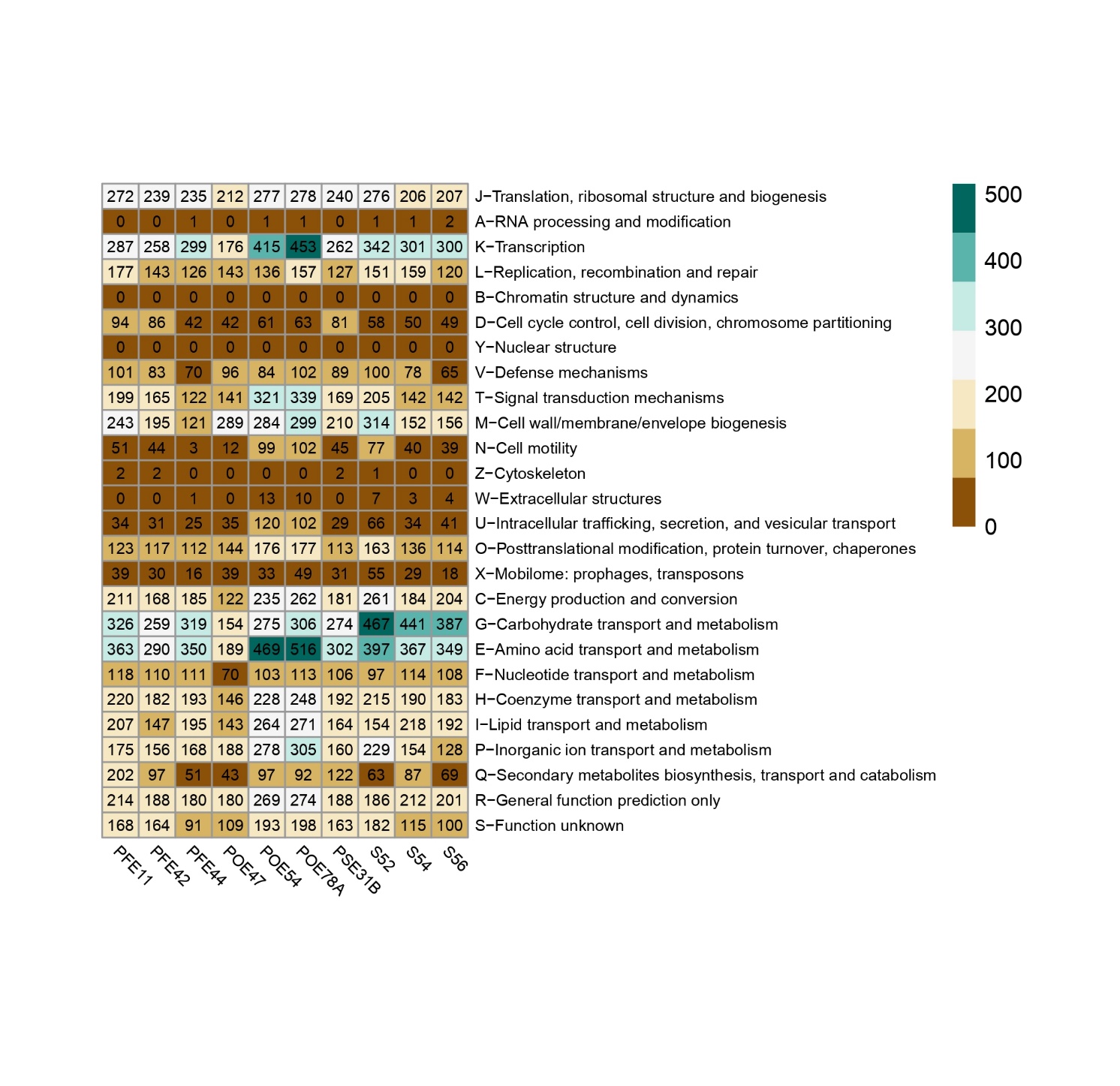

**Supplementary Figure 3.** Number of COGs for each genome with their related functions. PFE11, *Bacillus velezensis*; PFE42, *B. velezensis*; PFE44, *Glutamicibacter halophytocola*; POE47, *Chryseobacterium* sp.; POE54, *Pseudomonas salmasensis*; POE78A, *P. simiae*; PSE31B, *B. velezensis*; S52, *Leclercia* sp.; S54, *Paenarthrobacter ureafaciens*; S56, *Paenarthrobacter* sp.

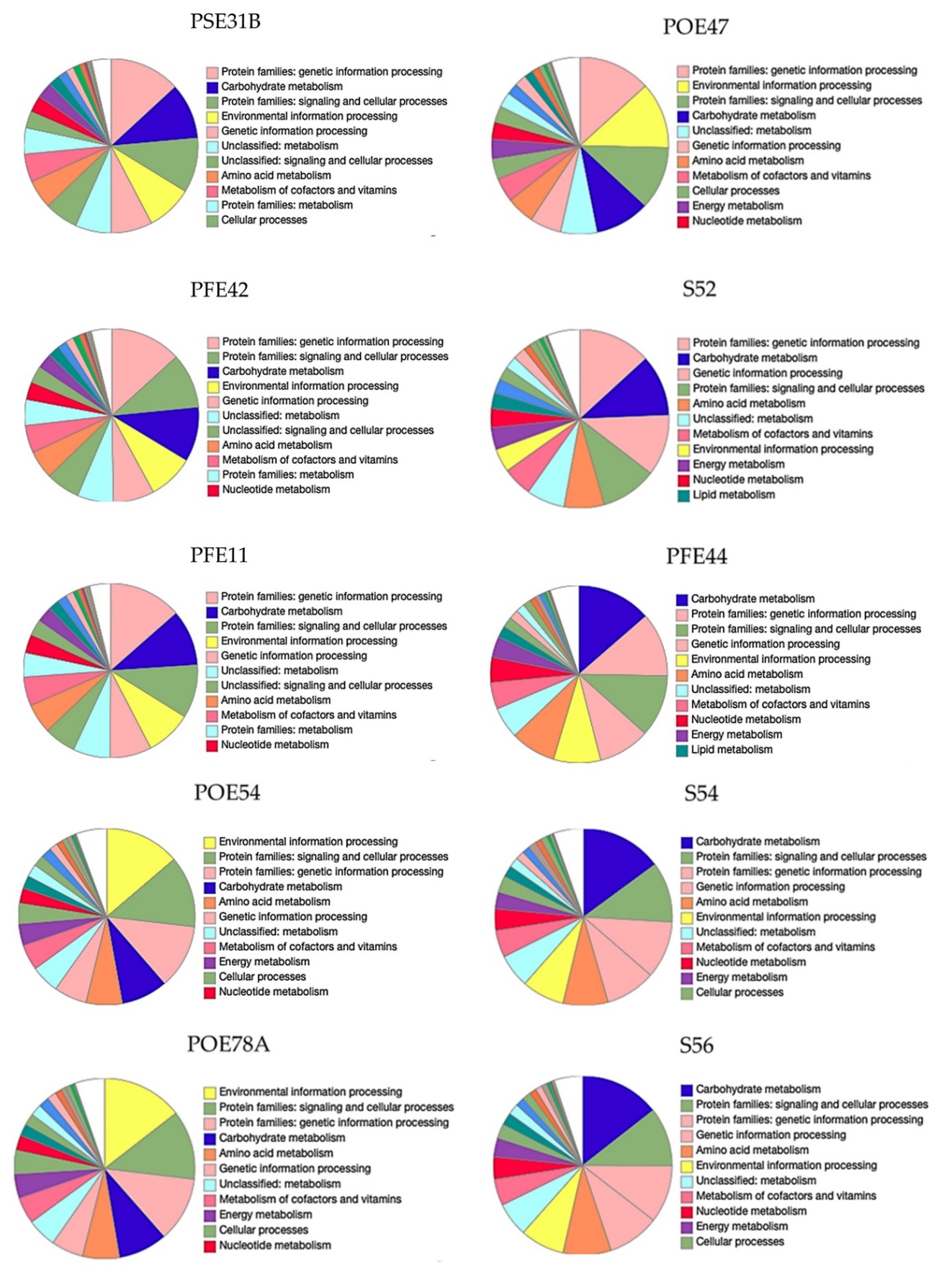

**Supplementary Figure 4.** Detailed distribution of KEGG in the ten genomes. PSE31B, *Bacillus velezensis*; PFE42, *B. velezensis*; PFE11, *B. velezensis*; POE54, *Pseudomonas salmasensis*; POE78A, *P. simiae*; POE47, *Chryseobacterium* sp.; S52, *Leclercia* sp.; PFE44, *Glutamicibacter halophytocola*; S54, *Paenarthrobacter ureafaciens*; S56, *Paenarthrobacter* sp.

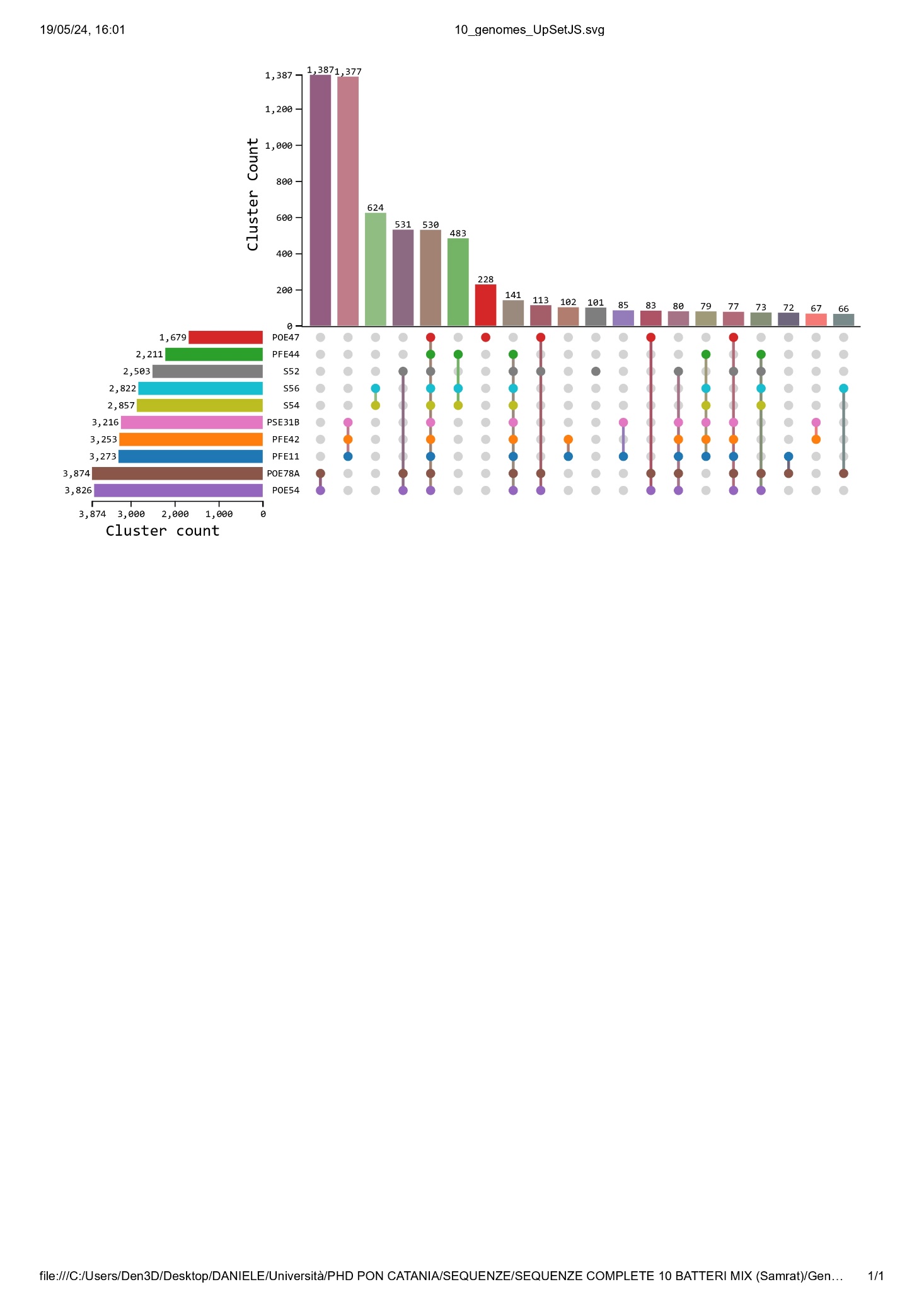

**Supplementary Figure 5.** Number of orthologous clusters in each species and shared orthologous gene clusters among species. Left horizontal bar chart shows the number of orthologous clusters per species, while the right vertical bar chart shows the number of orthologous clusters shared among the species. The lines represent intersecting sets. POE47, *Chryseobacterium* sp.; PFE44, *Glutamicibacter halophytocola*; S52, *Leclercia* sp.; S56, *Paenarthrobacter* sp.; S54, *P. ureafaciens*; PSE31B, *Bacillus velezensis*; PFE42, *B. velezensis*; PFE11, *B. velezensis*; POE78A, *Pseudomonas simiae*; POE54, *P. salmasensis*.

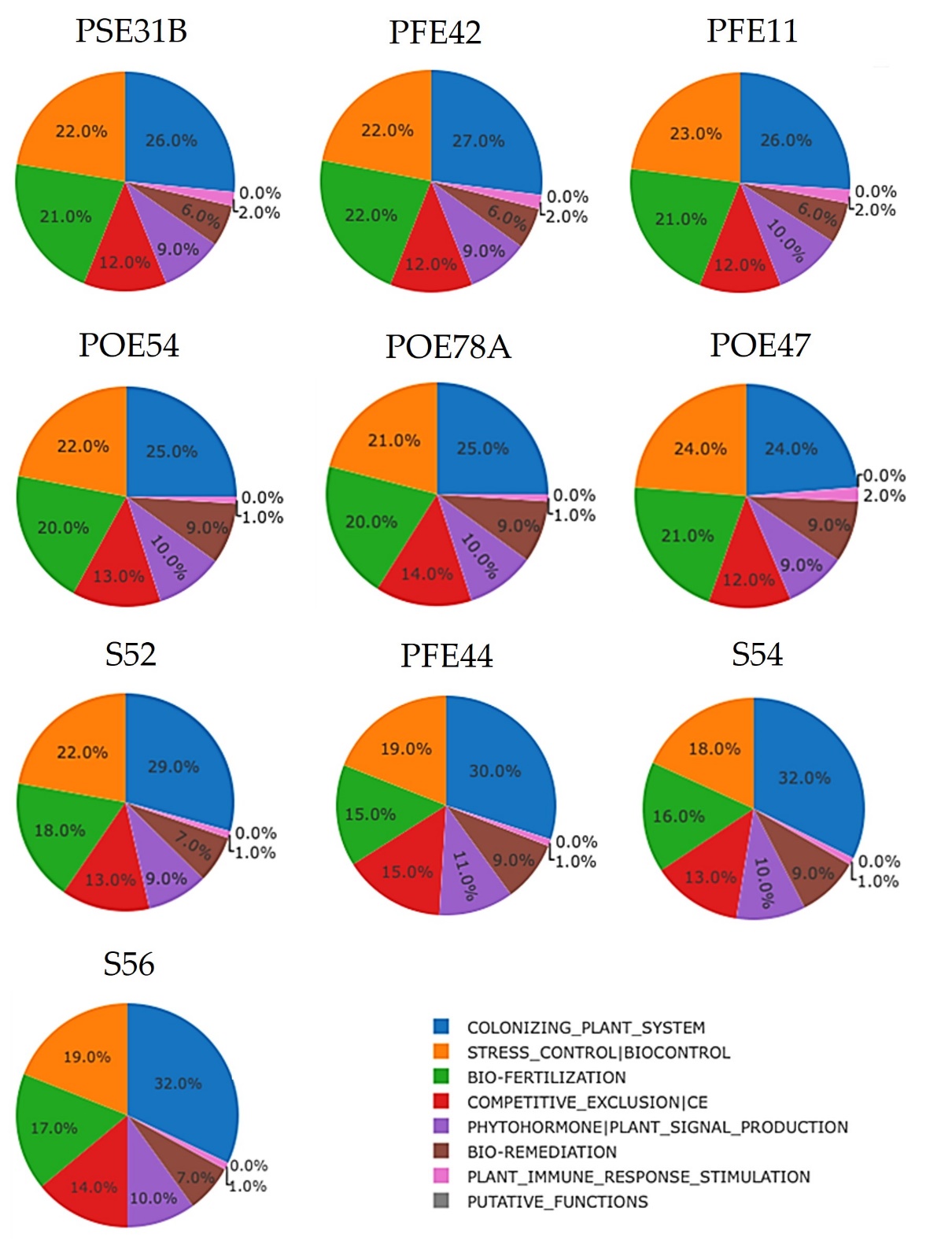

**Supplementary Figure 6.** Macro-categories of genes classes in the genomes of the ten strains. Plant growth promoting traits (PGPTs) were predicted using PGPT-Pred module of PLaBAse v1.01 (Patz et al., 2021). PSE31B, *Bacillus velezensis*; PFE42, *B. velezensis*; PFE11, *B. velezensis*; POE54, *Pseudomonas salmasensis*; POE78A, *P. simiae*; POE47, *Chryseobacterium* sp.; S52, *Leclercia* sp.; PFE44, *Glutamicibacter halophytocola*; S54, *Paenarthrobacter ureafaciens*; S56, *Paenarthrobacter* sp.
